# The impact of age on genetic risk for common diseases

**DOI:** 10.1101/2020.07.17.208280

**Authors:** Xilin Jiang, Chris Holmes, Gil McVean

**Author notes:** Corresponding author, Address: Big Data Institute, Li Ka Shing Centre for Health Information and Discovery, Old Road Campus, Oxford OX3 7LF, UK.

## Abstract

Inherited genetic variation contributes to individual risk for many complex diseases and is increasingly being used for predictive patient stratification. Recent work has shown that genetic factors are not equally relevant to human traits across age and other contexts, though the reasons for such variation are not clear. Here, we introduce methods to infer the form of the relationship between genetic risk for disease and age and to test whether all genetic risk factors behave similarly. We use a proportional hazards model within an interval-based censoring methodology to estimate age-varying individual variant contributions to genetic risk for 24 common diseases within the British ancestry subset of UK Biobank, applying a Bayesian clustering approach to group variants by their risk profile over age and permutation tests for age dependency and multiplicity of profiles. We find evidence for age-varying risk profiles in nine diseases, including hypertension, skin cancer, atherosclerotic heart disease, hypothyroidism and calculus of gallbladder, several of which show evidence, albeit weak, for multiple distinct profiles of genetic risk. The predominant pattern shows genetic risk factors having the greatest impact on risk of early disease, with a monotonic decrease over time, at least for the majority of variants although the magnitude and form of the decrease varies among diseases. We show that these patterns cannot be explained by a simple model involving the presence of unobserved covariates such as environmental factors. We discuss possible models that can explain our observations and the implications for genetic risk prediction.

**Author summary:** The genes we inherit from our parents influence our risk for almost all diseases, from cancer to severe infections. With the explosion of genomic technologies, we are now able to use an individual’s genome to make useful predictions about future disease risk. However, recent work has shown that the predictive value of genetic information varies by context, including age, sex and ethnicity. In this paper we introduce, validate and apply new statistical methods for investigating the relationship between age and genetic risk. These methods allow us to ask questions such as whether risk is constant over time, precisely how risk changes over time and whether all genetic risk factors have similar age profiles. By applying the methods to data from the UK Biobank, a prospective study of 500,000 people, we show that there is a tendency for genetic risk to decline with increasing age. We consider a series of possible explanations for the observation and conclude that there must be processes acting that we are currently unaware of, such as distinct phases of life in which genetic risk manifests itself, or interactions between genes and the environment.

## Introduction

Many studies have demonstrated the potential utility of using genetic risk factors for predicting individual risk of common diseases, ranging from heart disease (Ripatti et al., 2010; Rossouw, 2002) to breast cancer (Mavaddat et al., 2015) and auto-immune conditions (Cotsapas et al., 2011). Genetic risk coefficients can be estimated from cross-sectional genome-wide association studies, which estimate enrichment of common genetic variants among clinically-ascertained (or sometimes self-reported) cases. Genome-wide scores, typically referred to as polygenic risk scores (PRS), are usually constructed as linear combinations of individual variant effects, though there is considerable variation in how variants are selected for inclusion and how coefficients are estimated (Choi et al., 2020). Nevertheless, validation on independent data sets has demonstrated odds-ratios for PRSs that are comparable to established risk factors, both lifestyle-related (Mosley et al., 2020) and monogenic (Khera et al., 2018), thus providing an impetus for their adoption within health management, both at individual and population levels; though see (Mosley et al., 2020).

One aspect of genetic risk estimation that has received relatively little attention is the role of age in modulating effects. Several studies have identified variants that influence age-at-onset for diseases including type 1 diabetes (Ide et al., 2002), Alzheimer’s disease (Wollmer et al., 2003) and multiple-sclerosis (Moutsianas et al., 2015). Often, variants identified are the same as those affecting lifetime risk. Similarly, individuals with high PRS risk tend to have earlier age-at-onset than those who have low genetic risk, but nevertheless get the disease (Harbo et al., 2014; Nalls et al., 2015) and genetic analyses of quantitative traits including blood pressure, lipid levels and BMI have identified genetic variants whose effect size changes with age (Dumitrescu et al., 2011; Lasky-Su et al., 2008; Mostafavi et al., 2020; Shi et al., 2009; Simino et al., 2014). These results raise the possibility that genetic risk factors may play larger or smaller roles in influencing risk of disease during different age intervals. However, the longitudinal analysis of disease risk has to account appropriately for the selection bias that arises in age-stratified analyses; even under a time-invariant proportional hazards model, those entering the disease state earliest will tend to be those with the highest burden of risk factors. This can be particularly problematic when not all risk factors are measured, as hidden risk factors will act to apparently dilute risk over time, a phenomenon typically referred to as frailty in the epidemiological modelling literature (Aalen, 1988; Govindarajulu et al., 2011; Lin et al., 1998).

Here, we address two open questions in the analysis of longitudinal genetic risk for common disease. First, we introduce a method to infer the nature of the relationship between age and genetic risk for individual variants that is appropriate for censored data such as that available from biobanks. Because the information available for single variants is relatively weak, we use a Bayesian clustering approach to identify sets of variants that show similar profiles of risk with age. On applying the method to data from the UK Biobank on 24 common diseases, our primary finding is that, in agreement with previous observations, where we find evidence for non-uniform risk profiles, genetic factors most strongly influence risk of early disease. However the quantitative nature of the relationship between genetic risk and age varies among diseases and, for some, we find evidence for multiple, distinct profiles of age-varying risk. Second, we consider whether observed patterns can potentially be explained by the presence of unmeasured risk factors. To achieve this, we fit parametric models that accommodate unmeasured variation in risk to age-varying incidence and use these models to predict the drop-off in apparent genetic risk that could be attributed to frailty. We find that the observed drop-off in risk has a qualitatively different profile from that expected from simple models of frailty. Rather, our observations argue for greater biological complexity, such as genetic risk primarily affecting developmental phases in the evolution of individual liability or the presence of interactions among risk variants and environment.

## Results

### Data preparation

We used the genotype data, individual information and Hospital Episode Statistics (HES) data from 409,694 individuals of British Isles ancestry in the UK Biobank dataset (Bycroft et al., 2018). We identified 24 disease-specific ICD-10 codes (Table 1; Table S1) with a prevalence > 0.5% and for which at least 20 independent associated variants were identified using the TreeWAS model (Cortes et al., 2020). For each ICD-10 code, we combined the primary and secondary diagnosis from the full HES data set. We used the starting date of the first episode that records the disease diagnosis to compute age for first disease onset (the difference between onset date to the month of birth rounded to the nearest year). The age at observation endpoint is recorded as either the age of individuals at the last update of the data set (here 2018-02-14) or the age of death if a death event is recorded. We used eight age intervals of five years each.

### Age-profiles for genetic risk scores

To first demonstrate that age-varying genetic risk is a common feature of complex disease we estimated genetic risk coefficients through logistic modelling of a training case-control study (across all ages) and then assessed the efficacy of a combined genetic risk score to differentiate between cases and controls within each age group in an independent testing set (see Methods, Fig 1 and Fig S1). For many diseases, and notably those identified later as having statistically significant evidence for non-uniform genetic risk profiles, we found a typically decreasing risk profile. For example, the odds-ratio for the 90th percentile of GRS for I25.1 “atherosclerotic heart disease of native coronary artery” drops from 3.63 in the youngest age group to 1.77 in the eldest. We also note while some disorders, such as E78.0 “pure hypercholesterolemia”, show a very dramatic decrease in risk between the youngest age groups, others, such as a I10 “essential (primary) hypertension”, show a much more gradual decline. These results suggest that the relationship between genetic risk and age varies among diseases and may indeed vary among variants, and motivates a more principled approach to the analysis of such data.

**Fig 1.**
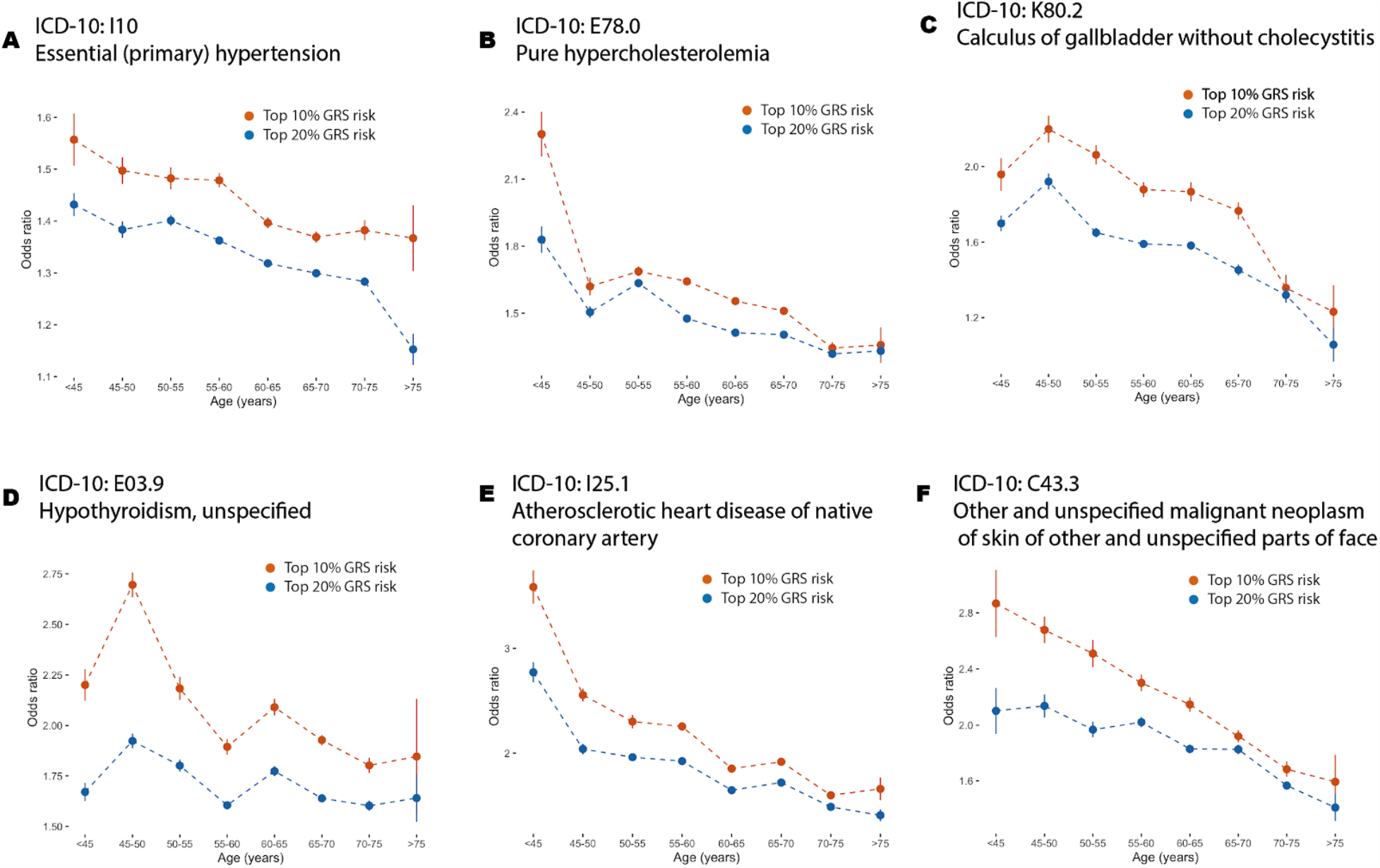
Age-varying genetic risk score (GRS) prediction power. A-F) Genetic risk score prediction power for six disorders where there is evidence for single non-constant profile, “Primary (essential) hypertension” (ICD-10 code I10), “pure hypercholesterolaemia” (E78.0); “Calculus of gallbladder without cholecystitis” (K80.2) and “Hypothyroidism, unspecified” (E03.9); “atherosclerotic heart disease of native coronary artery” (I25.1) and “other and unspecified malignant neoplasm of skin and unspecified parts of face” (C44.3). Curves for all diseases are shown in Fig S1. Odds ratios for the 80th (blue) and 90th percentiles of a combined genetic risk score within matched case-control samples (five controls for each case) are shown for each age interval; points indicate the average odds ratio of twenty five-fold cross-validation analyses with lines indicating the 95% confidence interval.

### Statistical inference of age-varying genetic risk with multiple variant categories

To estimate age-specific effects of variants we divided age into eight age intervals and used an interval-censoring approach in which the hazard rate for the risk factor is estimated by comparing those whose first disease event occurs during the interval in question to those who have a non-disease censoring event during the interval (such as death from a different disease, or drop-out from the study for reasons unrelated to disease) along with those who have neither a disease nor a censoring event during the interval (Fig 2). For a given variant, we estimated the effect size and its standard error for each interval using a proportional-hazards approach, matching additional covariates such as date of birth, sex, BMI and 40 genetic principal components (see Methods). Effect sizes for individual SNPs were estimated in both univariate and multivariate settings (see below). Because estimated variant-interval coefficients have high uncertainty, we used a Bayesian clustering approach to estimate latent profiles of age-specific genetic risk, encouraging smoothness of profiles through spline functions. Finally, to test for deviations from homogeneity of risk over age, and to test for the presence of multiple age-specific risk profiles, we used a permutation strategy. Full details of the methods are given in the Methods and Analytical Note.

**Fig 2.**
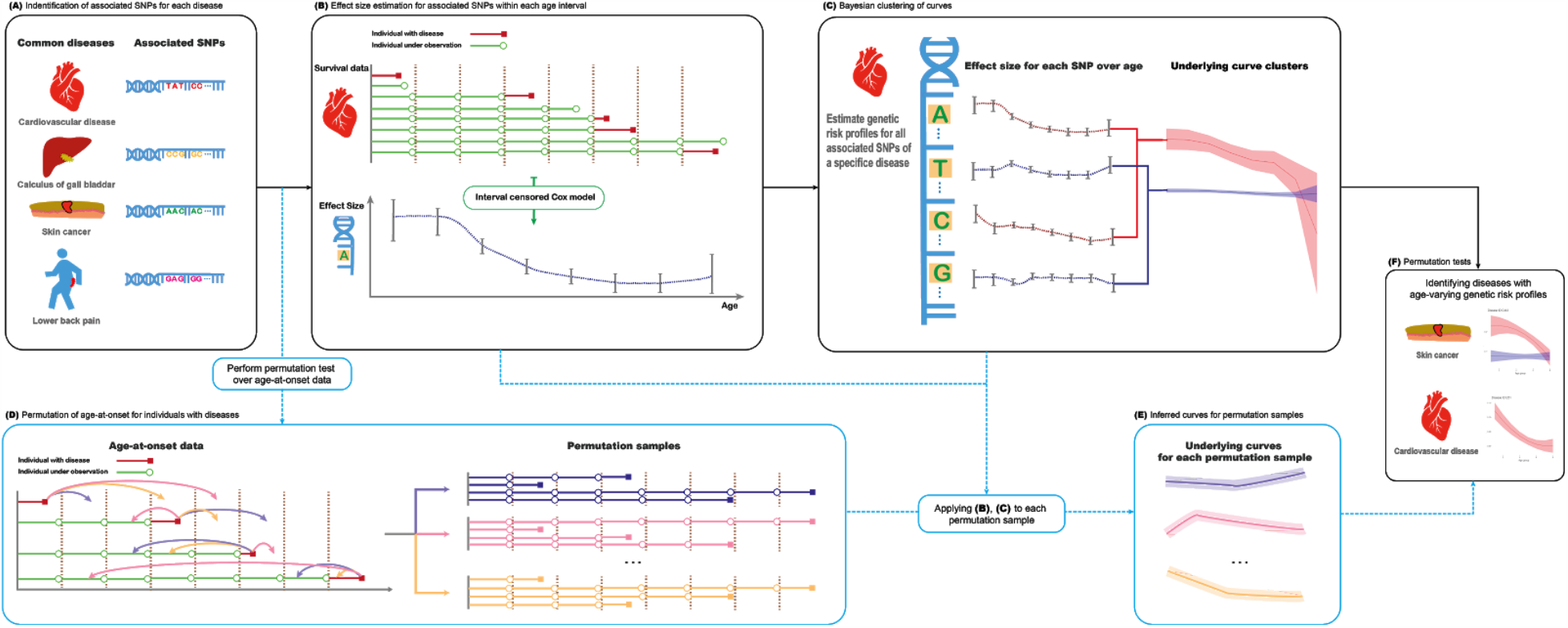
Schematic representation of methodology. A) Independent variants associated with a trait of interest are identified by analysis of the entire UK Biobank cohort using the TreeWAS methodology (Cortes et al., 2017). B) An interval-censored proportional hazards model (Finkelstein, 1986) is used to estimate the effect (and associated standard error) of each variant on the trait of interest within each of eight age intervals. C) Bayesian clustering is used to estimate age-profiles of risk, using either linear models or quadratic polynomials to encourage smoothness. D-F) Permutation is used to test for age-homogeneity of effect size as well as to assess the evidence for multiple age profiles.

To evaluate the methodology under the assumptions of the fitted model, we used stochastic simulation, varying the number of distinct profiles and their departure from uniformity. We first considered a likelihood ratio test (LRT) approach, fitting a linear model for risk profiles over age. Under realistic assumptions about the magnitude of effect sizes and number of associated variants we found that the multivariate approach is well-calibrated in its rejection of the null model of uniformity (i.e. when effect sizes are constant over time the LRT test has a false positive rate of 0.048 at P ≤ 0.05). When effect sizes are the same for all variants but these change by at least 0.6% per year (either increasing or decreasing), our approach has over 90% power to reject uniformity (Fig 3A). When quadratic polynomials were used to capture a wider range of possible risk profiles, we found that the LRT was less well calibrated under the null (false positive rate of 0.0725 at P ≤ 0.05; Fig 3A), hence we adopted a permutation strategy for analysing empirical data. When applying the quadratic model to data simulated under a linear profile, we find a good match between true and inferred profiles (Fig 3B).

**Fig 3.**
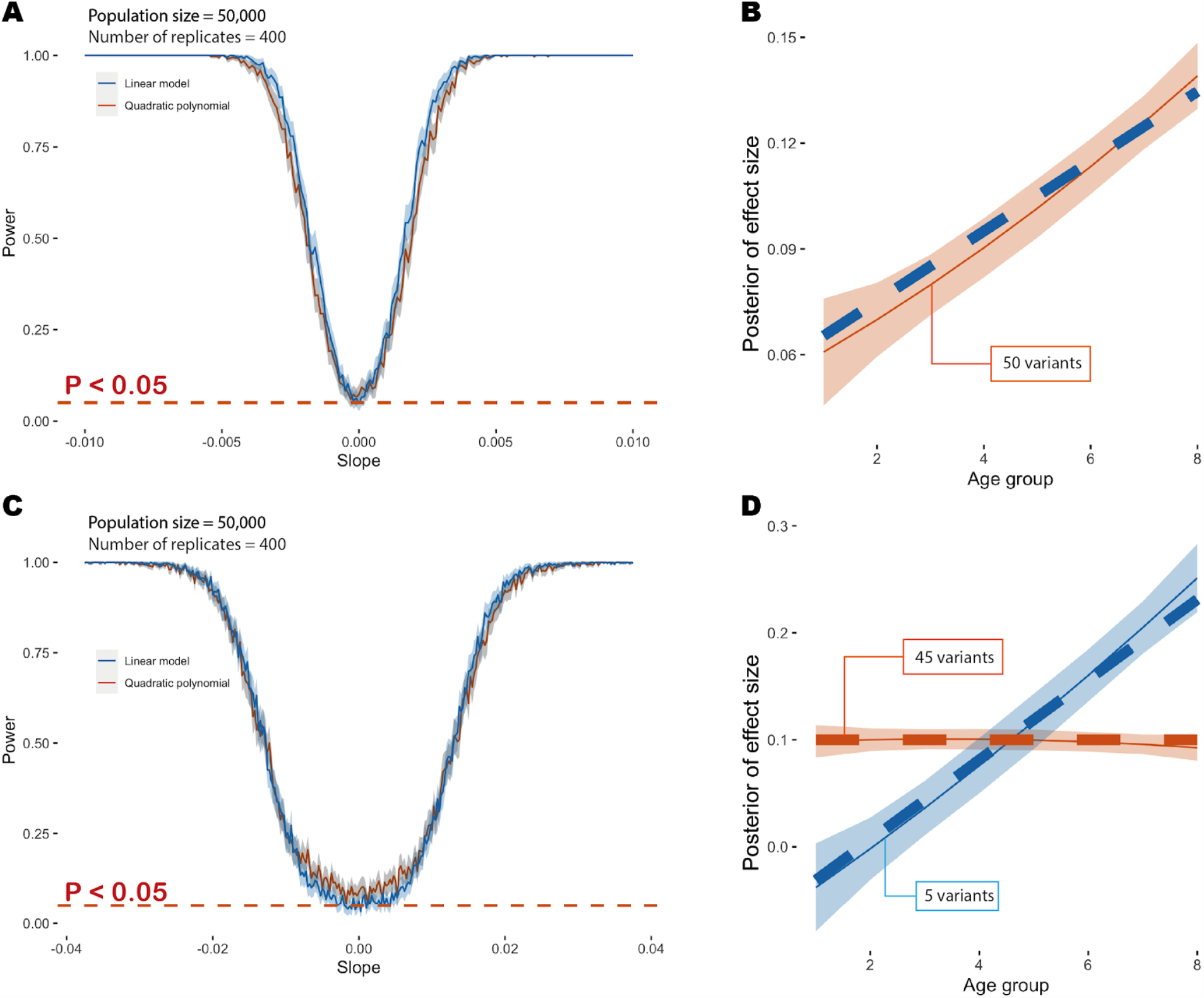
Overview of simulation results. A) Power at P ≤ 0.05 to detect deviation from age-homogeneity as a function of slope in a model where effect sizes change linearly with age. The blue line indicates the point estimate when using a linear model to fit, the red line indicates the point estimate with a quadratic polynomial model and the grey shading indicates the 95% confidence interval. B) Example showing the age-profile under which data are simulated (dashed blue line) and the inferred age profile (dashed red line) and 95% credible interval (red shading). C) Power at P ≤ 0.05 to detect multiple age profiles in a simulation where 90% of variants have a time-invariant profile and 10% have an effect size that increases with age. The solid blue line indicates power when fitting a linear model and the solid red line indicates power when fitting a quadratic model. The dashed red line indicates the nominal significance threshold. Note the change in x-axis scale compared to Fig. 2A. D) Example showing inferred age-profiles for the two components (mean posterior and 95% credible interval). Additional simulation details are provided in the Methods and Fig S2.

To simulate multiple cluster profiles, we modelled 10% of the variants as having a shared linear slope (the remainder being constant over age) and used a LRT to assess the evidence for multiple risk profiles. Here, we found that a 4% per year change in risk was required to achieve 90% power (at P ≤ 0.05) to detect multiple clusters (Fig 3C). Under the null (all variants have a constant profile) the test has a false positive rate of 0.063 for the linear and 0.088 for the quadratic polynomial fitting at P ≤ 0.05. When using the quadratic model to fit risk profiles we find a good match between true and inferred profiles (Fig 3D). We therefore conclude that the approach has sufficient power to detect deviations from constant profiles and provide unbiased estimates of risk profiles in data sets of comparable size and complexity to the UK Biobank. When analysing multiple diseases we used a FDR approach to correct for multiple testing.

### Application to common diseases in the UK Biobank

To formally consider evidence for a non-linear relationship between genetic risk and age for the 24 diseases in Table 1, we applied the novel methods outlined above. When effects for variants are estimated jointly and fitted to a linear latent profile, we identified, through permutation, nine diseases with evidence (P < 0.05) of a departure from uniform genetic risk over age (Table 1). These are: C44.3 “other and unspecified malignant neoplasm of skin of other and unspecified parts of face”; C44.5 “unspecified malignant neoplasm of skin of trunk”; E03.9 “hypothyroidism, unspecified”; E78.0 “pure hypercholesterolemia”; I10 “essential (primary) hypertension”; I20.9 “angina pectoris, unspecified”; I25.1 “atherosclerotic heart disease of native coronary artery”; I25.2 “Old myocardial infarction” and; K80.2 “calculus of gallbladder without cholecystitis”. All diseases have Q < 0.1 after FDR analysis. To model non-linearity we compared polynomial and cubic spline models with different degrees of freedom (Fig S3) and selected the quadratic polynomial model using likelihood ratio tests. No additional diseases were identified as having non-constant risk profiles when fitting a quadratic polynomial and only four of the original nine (E78.0, I10, I25.1 and C44.5) remain significant (Table 1). However, we find one additional disease (I20.0 “unstable angina”) and three of the above diseases (C44.3, E78.0 and I25.1) show evidence for more than one age-related risk profile (P < 0.05; Table 1, though only I25.1 has Q < 0.1).

As in the genetic risk score analysis, a common feature of the estimated risk profiles over age is a trend towards smaller effect sizes with increasing age (Fig 4A-D). For example, for I25.1, we find posterior of effect size drops by 50% from 45 years old to 75 years old and for C44.5 we find the posterior drops by 58% over the same interval. (Table S2). Where diseases may have multiple risk profiles (Fig 4E-F), at least one of these is also typically decreasing with age. Profiles for all 24 diseases are shown in Fig S4 and Fig S5. We find no compelling examples of increasing risk over age. These results are consistent with the effects of genetic risk factors to have a larger impact on the risk of early disease (de Miguel-Yanes et al., 2011; Mostafavi et al., 2020), rather than late disease, though it is important to note that the absolute rate of disease typically increases with age for all diseases studied here. Estimates of genetic risk profiles (under a model of one variant class) are provided in Table S3.

**Fig 4.**
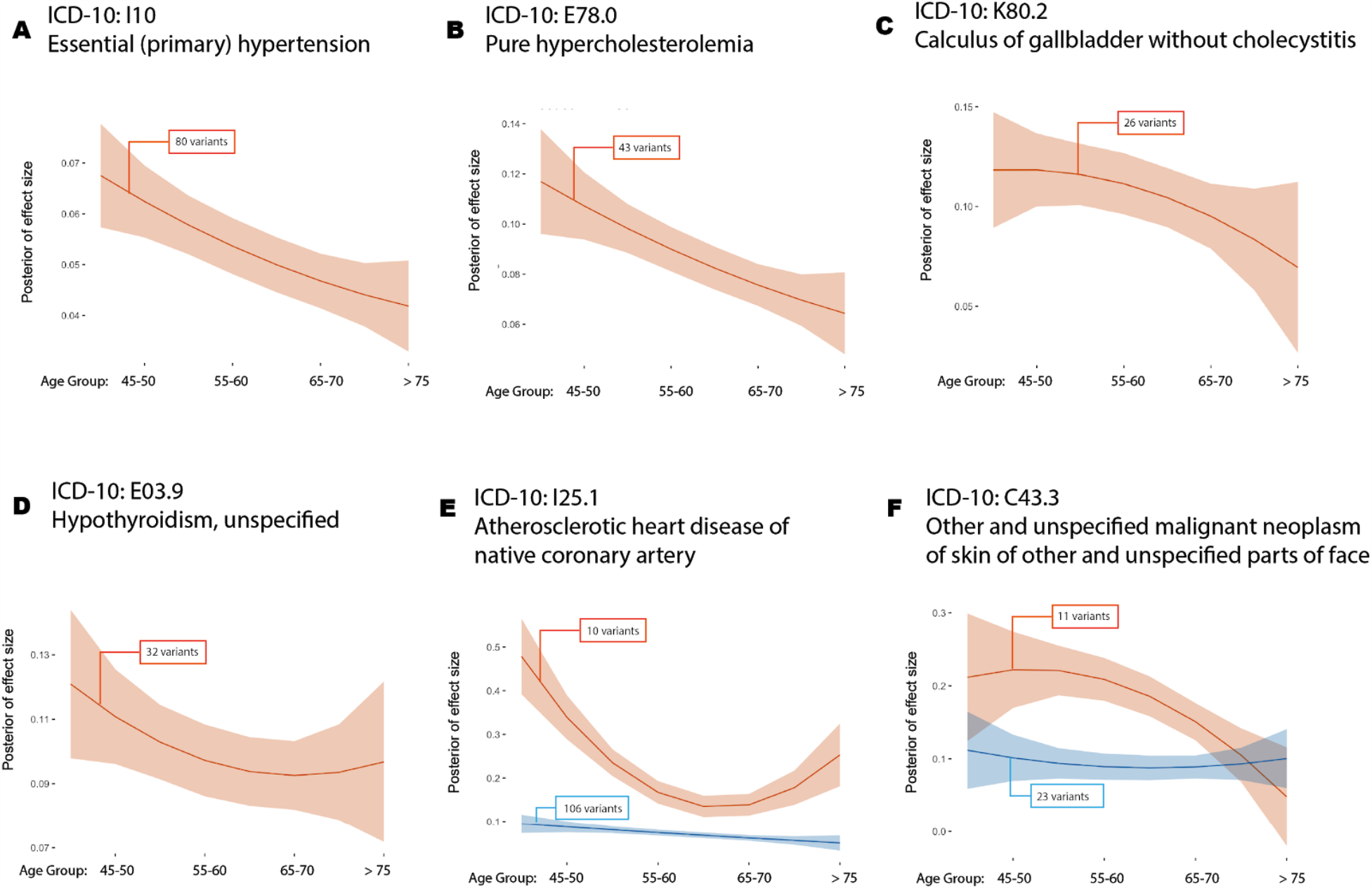
Age-varying disease risk profiles. A-D) Inferred cluster profiles for four disorders where there is evidence for single non-constant profile; “Primary (essential) hypertension” (ICD-10 code I10; P = 0.0001), “pure hypercholesterolaemia” (E78.0; P = 0.0001), “Calculus of gallbladder without cholecystitis” (K80.2; P = 0.0236) and “Hypothyroidism, unspecified” (E03.9, P = 0.0329); E-F) Inferred cluster profiles for two disorders where there is evidence for multiple non-constant profiles; “atherosclerotic heart disease of native coronary artery” (I25.1; P = 0.0001) and “other and unspecified malignant neoplasm of skin and unspecified parts of face” (C44.3; P = 0.0092). Curves for all diseases are shown in Fig S4. The solid line indicates the posterior mean and the shaded area the 95% credible interval; Numbers in parentheses indicate the number of variants in each cluster; All estimates are made with quadratic models for age-varying risk profiles.

### The impact of unobserved risk factors

One possible explanation for the decreasing impact of genetic risk is the presence of unobserved risk factors. For any causal covariate of interest, the presence of unmeasured and causally-associated uncorrelated covariates has the effect of generating (at the population level) additional variability in hazard rates, centred on the effect size. Such heterogeneity, historically referred to as frailty in epidemiology (Govindarajulu et al., 2011), has the potential to induce bias in effect sizes over time, somewhat remarkably even if independent of the covariate of interest, due to the increased rate at which individuals with high unmeasured risk enter into a disease state. Over time, those individuals with a risk-increasing covariate, but who do not have the disease, will become enriched for a protective background. Frailty will thus tend to lead to an underestimate of true effect sizes in older populations and, consequently, can even lead to biased effect size estimates (typically underestimates) in regression analysis of the entire cohort (Lin et al., 1998). To demonstrate the impact of such covariates we repeated the simulations under a constant risk profile, but multiplied individual risk by an unobserved factor that is generated from a gamma distribution (Fig S1A). We find that our test for age-dependence has a false positive rate of above 0.05 if the variance in risk is greater than 0.1 of the mean (Fig S1B).

To investigate the extent to which unmeasured genetic factors might be responsible for the diminishing of risk over time we first compared the results of univariate and multivariate analyses of the variants analysed here (Fig 5A). We found that results were essentially identical under the two approaches, suggesting that genetically-arising frailty cannot explain the pattern. We next attempted to estimate general parameters of frailty using incidence data from the UK Biobank by fitting a parametric model in which the underlying disease incidence (baseline hazard rate) increases in proportion to age as a power function of age, but where there is a distribution of rates within the population, parameterised as a gamma distribution with a mean of one and an unknown variance (Aalen, 1988; Vaupel et al., 1979); see Methods and Analytical Note. Estimates of parameters are provided in Table S4, along with the significance value for a goodness-of-fit test for the inferred model. We find substantial variation across diseases in the inferred parameters. For example, the baseline hazard rate of K80.2 “calculus of gallbladder without cholecystitis” is estimated to increase proportional to age to the power of 1.9, but with substantial frailty (scale parameter = 1.87, goodness-of-fit P = 0.93; Fig 5B). In contrast, the baseline hazard rate of C44.3 “other and unspecified malignant neoplasm of skin of other and unspecified parts of face” is estimated to increase more rapidly with age (power of 3.58), but with lower frailty (scale parameter = 0.94; P = 0.76). It should be noted that the simple parametric model can be rejected at P < 0.01 for only one (J45.9, “other and unspecified asthma”) of the 24 disorders, with the main discrepancy being a reduction in incidence among the eldest UK Biobank participants compared to the fitted model, which may potentially be explained by selection bias in recruitment and competing risks of multi-morbidity. We note that the estimated magnitude of frailty is typically sufficient to lead to an elevated false positive rate of the test.

**Fig 5.**
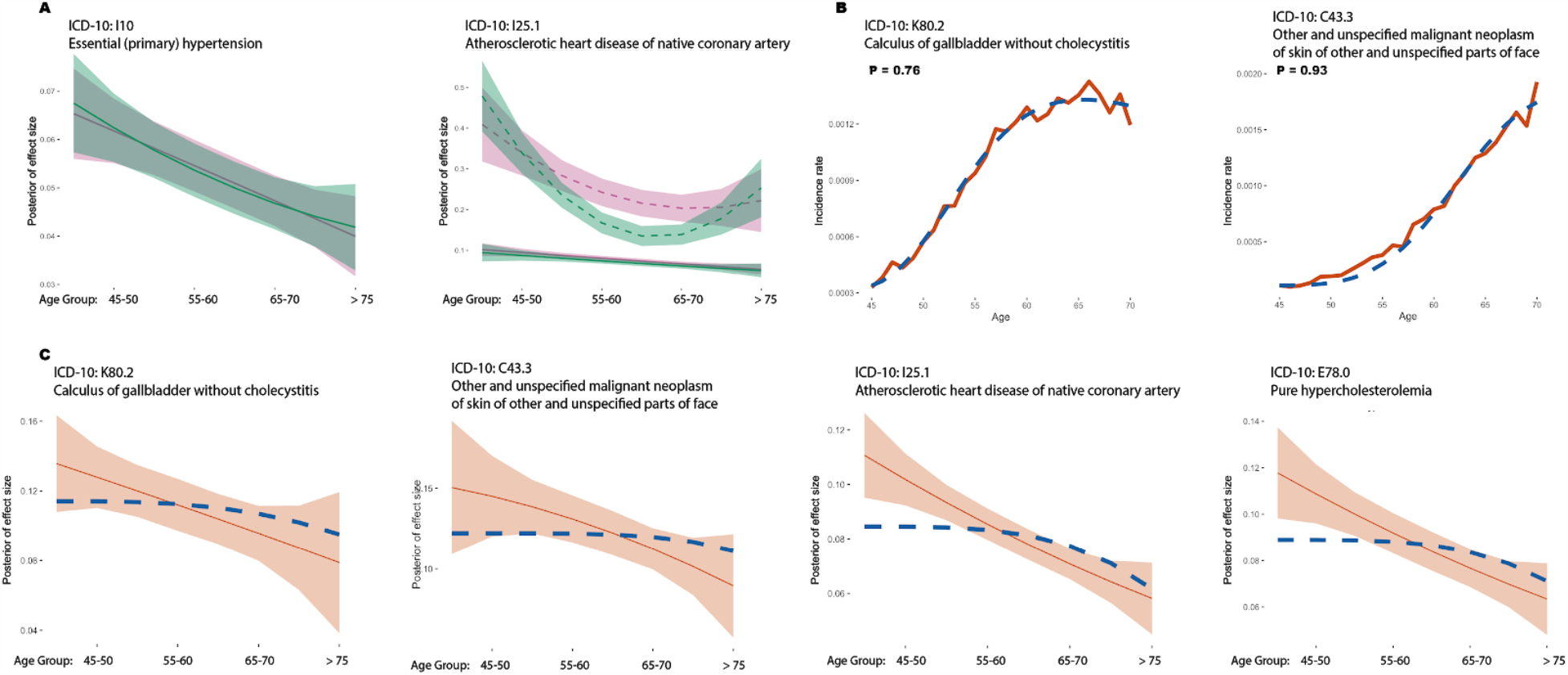
The impact of frailty on genetic risk profiles. A) Estimated age-profiles for genetic risk for I10 “essential (primary) hypertension” (left) and I25.1 “atherosclerotic heart disease of native coronary artery” (right) fitted under the univariate (purple) and multivariate (green) approaches. The solid line indicates the posterior mean and the shaded area the 95% credible interval. Comparisons for all diseases are shown in Fig S6. B) Estimated incidence by age for K80.2 “Calculus of gallbladder without cholecystitis” (left) and C44.3 “Other and unspecified malignant neoplasm of skin and unspecified parts of face” (right). The red solid line indicates the rate estimated from the UK Biobank (see Methods) and the dotted blue line indicates the fitted incidence curve from the parametric model. The P value indicates the Goodness-of-Fit test. Curves for all diseases are shown in Fig S7. C) Comparison of inferred genetic effect sizes (red curve) and those implied by the frailty parameters estimated from incidence rate within the UK Biobank (blue dashed curve).

Previous work has demonstrated that the magnitude of the diluting impact of frailty on effect sizes in longitudinal models can be predicted using the incidence and frailty distribution parameters (Aalen, 1988); notably the implied effect size at a given age is reduced by a factor proportional to the prevalence at that age multiplied by the variance of frailty distribution; see Methods. We therefore compared inferred (univariate) curves for genetic variants against that implied by the fitted frailty model (Fig 5C). In 17 of the 24 diseases we find that while the estimated frailty predicts a decreasing genetic effect size with age, the observed decrease both starts earlier and is of a larger magnitude than expected (Fig S8; Fig S9). Importantly, the estimated effect size tends to decrease substantially even when the prevalence of the disease is very low. We therefore conclude that, even after accounting for independent unmeasured factors that influence disease risk, genetic risk decreases with age.

## Discussion

Genetic factors influence lifetime risk for common and complex diseases through modulating a large number of molecular, cellular and tissue phenotypes, many of which are also likely to be affected by acute exposure and persistent environment (Bønnelykke & Ober, 2016; Corominas et al., 2014; Stranger et al., 2017). Despite such complexity, remarkable progress has been made in identifying factors, both genetic and non-genetic, that influence risk, each of which may only have a small effect, but which, in aggregate, have substantial and clinically relevant predictive value (Gandal et al., 2016; Jostins & Barrett, 2011; Manolio, 2013). To date, relatively little attention has been paid to the extent to which risk prediction can be improved by allowing genetic risk to be modulated by context, such as age, sex and environment, though see (Favé et al., 2018; Mühlenbruch et al., 2013). Here, we set out to measure how one specific aspect of individual context, namely age, has a modulating effect on genetic risk. For example, whether there are windows during which genetic risks are particularly relevant to disease and, conversely, other windows in which genetics plays a lesser role. The methods introduced here provide a flexible framework in which to address this question, as well as considering heterogeneity among diseases and classes of variants.

By applying the methods to data from the UK Biobank, we have identified four aspects of the relationship between age and genetic risk. First, we have shown that for many diseases, but certainly not all, there is statistical support for a non-constant relationship between age and the influence of genetic risk. Second, in such cases, genetic risk has the greatest effect at earlier ages, though the magnitude and form of the drop-off varies among diseases. This result agrees with and generalises earlier reports carried out using approaches that do not address the selection biases inherent in stratified analysis of longitudinal data. (Aalen, 1988; Mostafavi et al., 2020; Vaupel et al., 1979) Third, there is relatively little evidence for different groups of variants having substantially different relationships between age and risk; where we identify weak evidence for multiple classes, the differences are in terms of the magnitude of the downward slope. Fourth, the drop-off in risk with age cannot be ascribed to hidden variation in unmeasured risk factors, though as discussed below, this may reflect the presence of biological processes that are poorly captured through the widely-used proportional-hazards model. We note that the drop-off in impact of genetic risk factors does not mean that they are not relevant in predicting later disease, which is typically when most diseases occur. Rather, our results imply that the factor by which genetic risk factor increases risk above baseline for someone in their 40s may be exponentially higher than for an equivalent person in their late 70s (Table S5). For example, the factor by which being in the highest decile of genetic risk for I25.1 “atherosclerotic heart disease of native coronary artery” increases incidence over baseline between 45 to 50 years old is 6.62, compared to only 2.4 between 70 to 75 years old.

What biological processes could lead to a diminished influence of genetic risk over time? Genetic risk factors, unlike environmental ones, are present from birth, while non-genetic risk factors tend to accumulate and evolve over time. Such a difference could lead to a reduced impact of genetics over time if genetic risk primarily influences developmental pathways, while non-genetic risk affects separate and later-impacting pathways, such as those involved in adult homeostasis (Fig 6A). However, there are several contexts where genetic and non-genetic risk are, at least in part, mediated by the same factor, such as the impact of LDL cholesterol on cardiovascular disease. In such cases, statistical interactions between genetic factors and the environment (or potentially among genetic risk factors) could have a diluting effect on genetic risk in a manner similar to frailty (Fig 6B). Here, an interaction means that the combined effects of the genetic and non-genetic risk factors is worse than expected from their independent contributions. Biologically, such an interaction could arise from threshold models of homeostasis (meaning the system can buffer only up to a certain level of challenge), though many other biological processes could potentially lead to statistical interactions at the population level. Proportional hazards models, which assume that risk factors multiply an underlying hazard rate, cannot capture such time-varying influences. Generalised risk processes such as threshold models (Duggirala et al., 1997) provide a potentially richer framework for modelling such effects.

**Fig 6.**
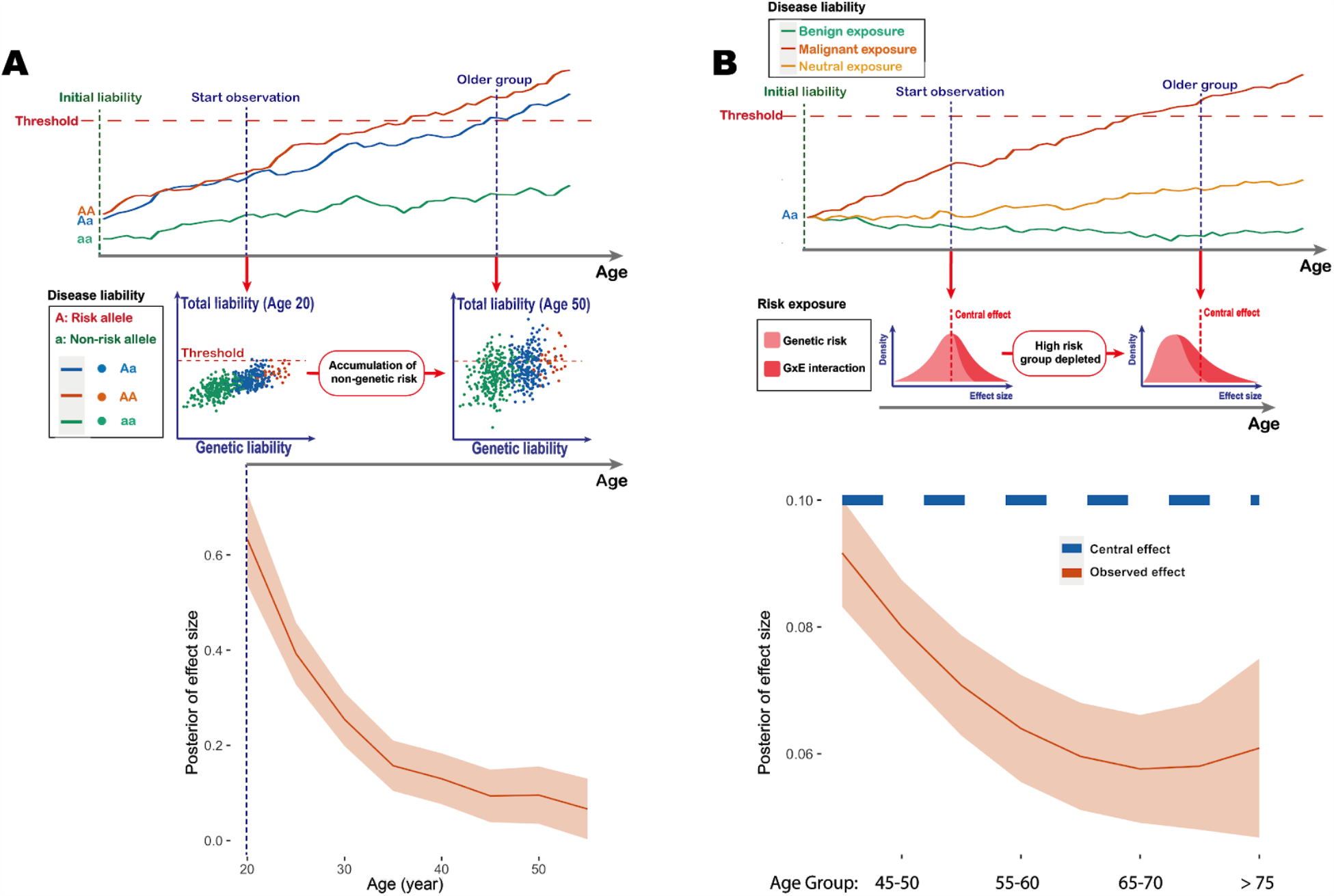
Models for a decreasing influence of genetic risk with age. A) A threshold model, in which each individual has a disease “liability” which evolves over age. Disease onset occurs when liability crosses a threshold. The upper panel shows example trajectories, where genetic risk alters only the liability baseline. The middle panel is a schematic representation of a simulation in which genetic risk affects developmental pathways at birth, while non-genetic risk accumulates over time. The lower panel shows an estimation of the effect size from a simulated dataset of UK Biobank sample size (see Methods). B) Interactions between genetic and environmental risk factors can create a distribution of effect sizes for a specific genotype. The upper panel shows example trajectories, where the environment influences the slope of the trajectory. The middle panel shows how individuals at higher risk enter disease earlier, diluting the effect size estimation at a later age. The lower panel shows simulation results under such a model using realistic parameters from UK Biobank (see Methods).

Whatever the cause of age-varying genetic risk, our results have several implications for the use of genetic risk factors in the genetic analysis and prediction of disease risk. Most obviously, genetic risk prediction for early disease is likely to be more effective than for later disease. For most of the diseases studied here, the inference of a single age-profile does mean that the rank order of genetic risk for an individual is stable over time. However, it implies that integrated prediction from genetic and non-genetic risk factors (Aschard et al., 2012; Kraft et al., 2007; Thomas, 2010) will have to consider the evolving contribution of genetics over age. For diseases with multiple age profiles, even the rank order of genetic risk among individuals could change over time. Finally, because contexts beyond age, such as sex and environment, modulate genetic risk (Mostafavi et al., 2020; Ober et al., 2008; Thomas, 2010), each of these will induce its own age-specific profiles. As a consequence, effective genetic prediction will most likely be driven by empirical models that can benefit from access to large and well-measured populations, such as population-scale biobanks.

## Materials and Methods

Full technical details are given in the Methods and Analytical Note.

### Data preparation

We use the genotype data, basic demographic data and Hospital Episode Statistics (HES) data from 409,694 individuals of British Isles ancestry in the UK Biobank dataset (Bycroft et al., 2018). 31 ICD-10 codes were identified with a prevalence above 5% and for which at least 20 independent associated variants were previously identified using the TreeWAS model (Cortes et al., 2020). Of these, we analysed 24 that correspond to specific disease conditions (as opposed to procedures) and that have sex- and age-distributions compatible with our framework. These are listed in Table 1. For each ICD-10 code, we combined the primary and secondary diagnoses from the HES data. We used the starting date of the first episode that records the disease diagnosis to define the age of disease onset, which is calculated as the difference between onset date to the month of birth (due to data privacy, we only have access to birth information specified to year and month). The onset age is rounded to whole years. For each ICD-10 code, only the first recorded diagnosis of each individual was used. For the population under observation, we also computed the age at observation endpoints, which is either the age of individuals at the last update of the data set (here 2018-02-14) or the age of death if a death event is recorded. We categorised the disease onset age into 8 age intervals, the first and last of which are “before 45 years old” and “after 75 years old”, with 5-year intervals in between.

We then constructed interval censored data sets for the selected disease. Each age interval is an observation window of all healthy (alive and without onset of target disease) individuals who survived past the starting point of the interval. Onset of disease and exiting the study (death or no further records available) are recorded as “case” and “censored” events respectively. Events happening after this interval are considered right-censored at the end of the interval. We then performed case-control matching over the sub-population observed within each interval in two steps. First, we divided the sub-population into a disease group and a control group. The disease group are those who have disease onset within the interval, and the control set are those who do not, including individuals who have disease onset after the age interval, so long as they remain healthy before the endpoint of the interval. This is what the term “interval-censored” means. In survival analysis, the control groups are called “censored”, and the age at the “censoring” event is also needed for unbiased estimation. If a censoring event is observed within the age interval (i.e. the age of a death record or the last update in the UK Biobank is before the age interval end point), we used the age at the censoring event. If an individual does not have an event record within the age interval, we take their age at the end of the interval, regardless of their future events. Second, for each case in the disease group, we pick four nearest neighbors (without replacement) from the control group, matching sex, BMI, year of birth and 40 genetic principle components. The covariates are available within the UK Biobank data set, over which we computed the principle components across the British Isles ancestry population. We then compute the Euclidean distance of the principle components to find the nearest neighbors in the population.

### Estimating age-dependency of genetic risk score in prediction

The SNPs of interest are obtained through prior TreeWas analysis, where we select variants that have Bayes factors (BF) ≥ 5 (BF is computed for a single variant’s effect over the TreeWas model) and posterior probability (PP) ≥ 0.99 for target diseases (Cortes et al., 2017). We further filtered the set of SNPs to ensure LD-independence (loci kept with absolute Pearson correlation coefficient smaller than 0.2).

To assess whether the collective effect of risk variants, as captured by a combined genetic risk score (or polygenic risk score - PRS), showed profiles of age-varying risk, we used the case-control matching procedure described above with five-fold cross-validation, keeping 20% of case-control pairs for each age interval as test sets, and estimating effect sizes for the selected variants in the remaining 80% of case-control pairs using multivariate logistic regression (including age, sex and 40 genetic PCs). The effect of the PRS on risk within each age interval in the test set was then estimated (again with logistic regression and covariates). We estimated the odds-ratio for the top decile of risk and the top 20% of risk, using 20 repeats of the procedure to obtain the bootstrap sampling distribution.

### Estimating age-specific effects of genetic risk factors

We used a standard GWAS approach to identify the risk and protective alleles at each locus (Chang et al., 2015), over the case-control matched dataset described above. The first 40 genetic principle components are taken as covariates. For all loci that have protective minor alleles (odds ratio < 1), we switched allele labels to assign consistency of risk direction.

To obtain an unbiased estimate of genetic risk effect size over age, we used a proportional hazard (PH) model to estimate the genetic hazard ratio for different age groups, using the case-control matched data set. Within each of the 8 age intervals, we applied the PH model to the disease group and control group, accounting the censoring effect. We used a univariate model to estimate the effect size of each variant separately and a multivariate model to estimate effect sizes for all variants jointly. Covariates include the first 40 genetics PCs of the UK Biobank and are regressed out for each interval. Both the point estimate and standard error of effect sizes are obtained for each variant within each age interval. These summary statistics are used subsequently for curve-cluster fitting.

### Bayesian clustering of genetic risk profiles

To group variants that have similar age-dependency, we applied a Bayesian clustering of curves model described (see Analytical Note). The model assumes each variant has an age-dependent effect profile which is generated from a mixture of curves model. The mean and standard error of the age-specific effects for individual variants described above are the inputs of the model, from which we infer the underlying generative latent curve. The model allows vertical translation in the generative process (i.e. the likelihood won’t change much if the profile of the variant is far from the latent profile, as long as the shape of the variant curve is similar). The latent curve is a spline whose smoothness is controlled by changing the degrees of freedom. For detailed specification of model and hyper-parameters, see the Analytical Note. Inference is performed by an EM algorithm and was repeated 20 times with random initialization of variables (see Analytical Note). The highest likelihood sequence was retained. Since EM only provides a point estimate, we estimate the curve’s credible interval using a variational approach. The inferred profiles with 95% credible intervals are shown in Fig S4. The derivation and proof of the approach are provided in Analytical Note.

### Permutation testing for genetic effect heterogeneity over age

To provide robustness in testing for age heterogeneity, we carried out a permutation test, using the likelihood ratio test statistic (for fitting a non-uniform genetic risk over age) for both the original data set and permutation samples to obtain the permutation p-value. The likelihood ratio test compares an alternative model with linear and quadratic genetic risk over age and a null model assuming a constant effect over age (see Methods and Analytical Note).

To perform permutation tests, we kept the matched case-control structure and then sampled case-control pairs for each age interval, while fixing the onset age distribution for permutation samples. We repeated the procedure 10,000 times to obtain permutation samples, and computed the likelihood ratio for each sample. We note that the likelihood ratio does not include the prior term for spline coefficients, while EM finds the Maximum a Posteriori (MAP) estimate (see Analytical Note), which will give a likelihood slightly lower than the MLE estimate. Under the permutation test framework, the p-values will be consistent as long as the same test statistics are used for both the original data set and permutation samples (Neyman et al., 1933). We further checked that the difference between MAP estimation and MLE estimation is negligible.

The EM procedure (see Analytical Note) is initialised randomly 20 times for the observed data and each permutation sample to compute the likelihood ratios. The permutation p-value for each disease is obtained from the likelihood ratios. We correct for multiple testing using FDR, with the corrected q-values shown in Table 1 (when a multivariate approach is used to estimate effect size) and Table S1 (when a univariate approach is used).

In order to determine the optimal number of clusters for each disease, we performed permutation tests using the same procedure, considering the addition of each new cluster. For adding the k+1 cluster, the alternative model has k+1 clusters and the null model has k clusters. All models are fitted with quadratic polynomials (see Analytical Note). Again, we computed the likelihood ratio statistics for both the observed data set and permutation samples to obtain p-values. This analysis is performed over all diseases and adjusted for multiple testing with FDR. We note that we found no compelling evidence supporting a model of more than two clusters for any disease. The p-values and q-values for the test of two clusters are shown in Table 1 (when effects for individual variants are estimated are estimated jointly) and Table S1 (when effects for individual variants are estimated using a univariate model).

### Estimating effects of unobserved risk background

To estimate the effect of unobserved risk factors, we assumed an individual hazard model that has a frailty coefficient and baseline hazard. The frailty coefficient has mean 1 and a scale parameter that controls the variance of population hazard rate. We chose the baseline hazard to be a power function of age. We fitted the model to the empirical incidence rates in the UK Biobank. The empirical incidence rate at a specific age is computed as the number of individuals who have first onset of the target disease within this age year, divided by the number of healthy individuals at risk at the beginning of this age year. We then fitted the parametric hazard to the empirical incidence rate until age 70, and finally subtracted the intercept from the empirical incidence rate to match the parametric form of the hazard rate. We fitted the model by minimizing square error using the Nelder-Mead method. The fitted incidence curves are compared with empirical curves for all diseases (Fig S7). We also computed a Goodness-of-fit p-value for each disease, comparing the match between fitted and empirical three-year incidence rates using a Chi-square test statistic. The Goodness-of-fit p-values are shown in Table S4. We used the inferred parameters to predict how genetic effects are expected to be diluted by the presence of frailty (Fig S9; for technical details, see Methods and Analytical Note).

### Simulation

In our simulation, we generate a risk profile over age for each variant from underlying curves with different slopes. The individual risk is then computed at different ages, which are then used to generate disease incidence events over the simulated population. We choose the population size to be 50,000, which is comparable to our empirical case-matching population size (the set of common diseases analysed each have ~10,000 cases in the UK Biobank and we match each case with four controls). We simulated 50 SNPs (MAF of each SNPs are sampled from uniform distribution 0-0.5). The risk effect for each SNP is sampled from a profile which changes linearly with age. The individual hazard within a specific age interval is computed as the exponential of genetic risk multiplied by a linearly increasing baseline hazard ratio.

For each interval, we simulated the time to the next event using a homogeneous Poisson process with the defined individual hazard rate. An individual with no event in this interval is considered as observed (censored). We record only the first event as the onset of the disease. The simulation is performed over a 40 year duration divided into eight 5-year intervals, as most of the disease onset occurs between ages 40-80 years old in the UK Biobank. In order to represent the end of observation (study drop-out) or death events in the cohort, a competing censoring process is sampled using a Poisson process of constant rate. The dropout/death and disease onset events are combined and we keep the first event, labelling it as either disease or censoring. For parameters setting in the simulation, see Methods.

To test our statistical model for inferring age-varying genetic effects, we simulated a population using the scheme described above and analysed it using the methods described above to infer the genetic risk profiles over age and the underlying curves that generated them. We simulated the cohort with different values of the slope, which represent different age dependencies, and tested whether our method could recover the simulated values. We then assessed the power of the statistical test to detect age-varying genetic effects. We simulated the genetic risk profile with the slope ranging from −0.01 (linearly decreasing with age) to 0.01 (linearly increasing with age), with a step size of 0.0001. The simulated population is analysed using the null model of a constant effect with age, and an alternative model of either a linear model, or a quadratic polynomial curve. A likelihood ratio test is performed to calculate the p-value, and we calculate the power of rejecting the null at a threshold of p = 0.05. For each slope, the simulation was repeated for 400 times to estimate the power and its standard error.

To test our statistical model for detecting multiple clusters of genetic risk profiles, we simulated disease cohorts with five (10%) of the variants that had effect sizes generated from a non-constant latent profile, while the effect sizes for the remaining 45 variants had a constant (age-invariant) effect. We assessed our model as to whether it can detect the presence of multiple clusters. The simulated cohorts are analysed with both a null model of a single quadratic polynomial curve, as well as the alternative model of two quadratic polynomial curves. For each simulation, we compute the p-value for the likelihood ratio test comparing two clusters against one cluster, measuring power at p = 0.05. We varied the slope of the non-constant profile to test how different the curve needs to be from a constant effect to be distinguishable by our model. Power is computed for slopes ranging between −0.0375 and 0.0375, with a step size of 0.00025. For each slope, the simulation was repeated for 400 times to estimate the power and its standard error.

To model possible mechanisms for the observed decline in genetic risk with age we simulated a threshold model in which each individual has an unknown “liability”, which evolved over time (Demenais, 1991). For a specific disease, onset occurs when an individual’s “liability” passes a certain threshold. We simulated a liability model for 50,000 individuals with a single genetic effect that alters the starting point of liability. Genotypes were simulated with a risk allele frequency 0.3. The liability is simulated as a stochastic process with starting points altered by genotypes. We then simulated increments of liability from a Gaussian distribution which controls the drift and variance of the stochastic process. The stochastic process models the disease risk increase over age through the drift, and the correlation of increments induced by the variance of Gaussian distribution creates a “momentum” such that an individual’s health status tends to improve or deteriorate over years at similar rate. We simulated for 60 years and considered an individual to have an onset of a disease when the liability (arbitrarily) reaches 0. We then estimated the effect size of the risk allele over the age interval 21-60. For parameters setting in simulating the stochastic processes, see Methods.

To consider whether the decreasing pattern could be explained by interactions (either gene-by-environment or gene-by-gene) we performed additional simulations. We modelled the interaction of a focal genetic effect with other unobserved risk factors. Assuming the effect size interacts with environmental or other genetic factors, the effect size for each individual is generated from a positively defined distribution. We can show that the estimated marginal effect size will be increasingly underestimated as age increases for all positive defined probability distributions (see Analytical Note). We then performed a simulation using the parameter settings described at the beginning of this section, but sampled an effect size for each individual from a gamma distribution. The effect size for each individual remains constant over age intervals. We then inferred the posterior of effect size, presented in Fig 6B. We note that this model is a generalisation of the concept of frailty in which one allele has greater frailty than the other.

## Supporting information

Analytical Note

## Description of Supplemental Data

Supplemental Data include nine figures, five tables, Methods and Analytical Note.

## Declaration of Interests

G.M. is a director of and shareholder in Genomics PLC, and is a partner in Peptide Groove LLP. The other authors declare no competing financial interests.

### Acknowledgements

This research has been conducted using the UK Biobank Resource; application number 12788. This work uses data provided by patients and collected by the NHS as part of their care and Support. Funded by Wellcome (BST00080-H503.01 to XJ, 100956/Z/13/Z to GM); the Li Ka Shing Foundation (to GM); The Alan Turing Institute, Health Data Research UK, the Medical Research Council UK, the Engineering and Physical Sciences Research Council (EPSRC) through the Bayes4Health programme Grant EP/R018561/1, and AI for Science and Government UK Research and Innovation (UKRI) (to CH). Computation used the Oxford Biomedical Research Computing (BMRC) facility, a joint development between the Wellcome Centre for Human Genetics and the Big Data Institute supported by Health Data Research UK and the NIHR Oxford Biomedical Research Centre. The views expressed are those of the authors and not necessarily those of the NHS, the NIHR or the Department of Health. We thank Brieuc Lehmann and Luke Jostins-Dean for discussion and comments on the manuscript.

## Data and Code Availability

The code generated during this study is available at https://github.com/Xilin-Jiang/longitudinal_genetic_analysis

