## Supplemental Figures and Tables for "The impact of age on genetic risk for common diseases"

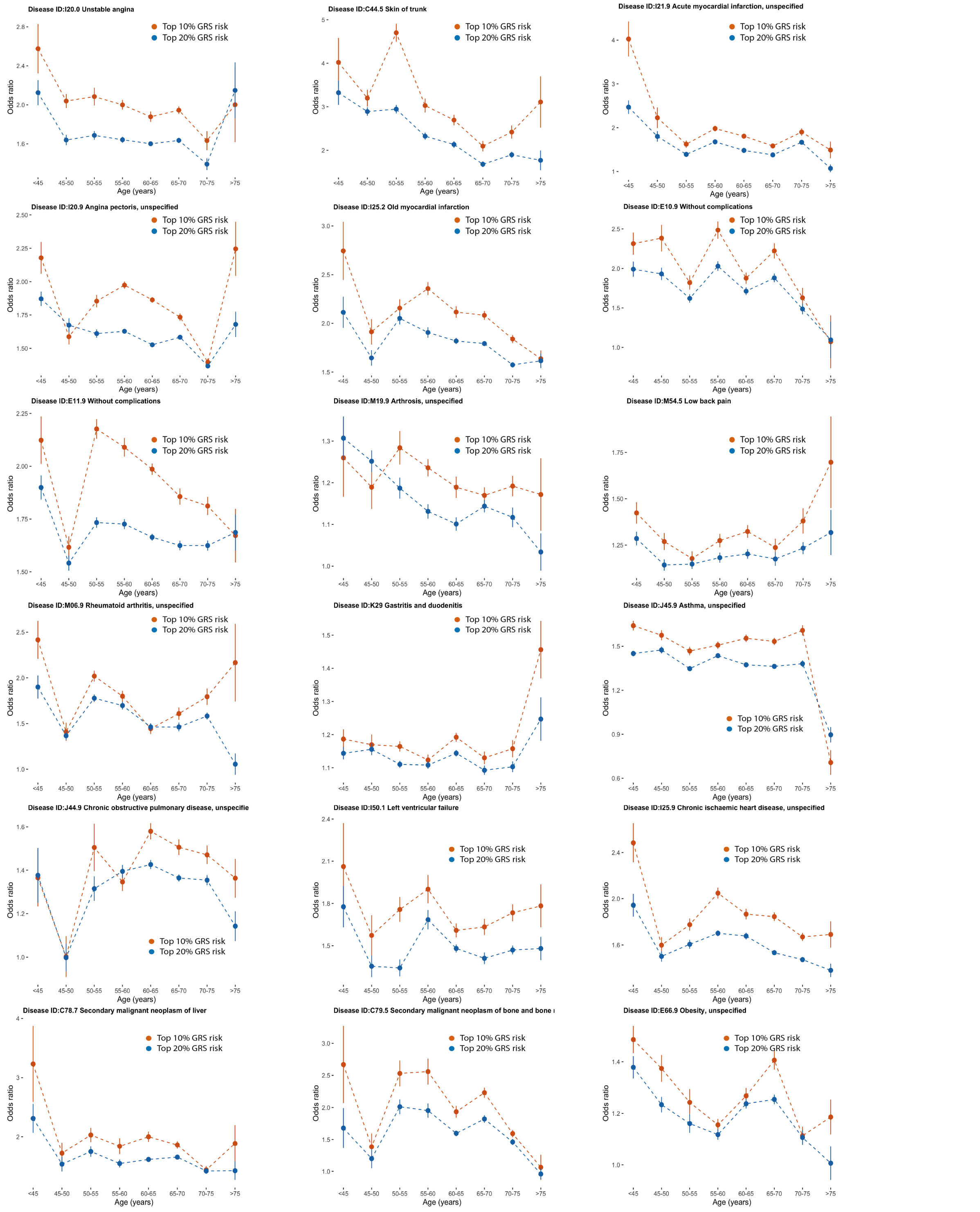


**Fig S1**: **The prediction power for combined genetic risk scores for additional diseases.** In each plot the odds ratios for the 80th (blue) and 90th percentiles of a combined genetic risk score within matched case-control samples (four controls for each case) are shown for each age interval; points indicate the average odds ratio of twenty five-fold cross-validation analyses with lines indicating the 95% confidence interval.

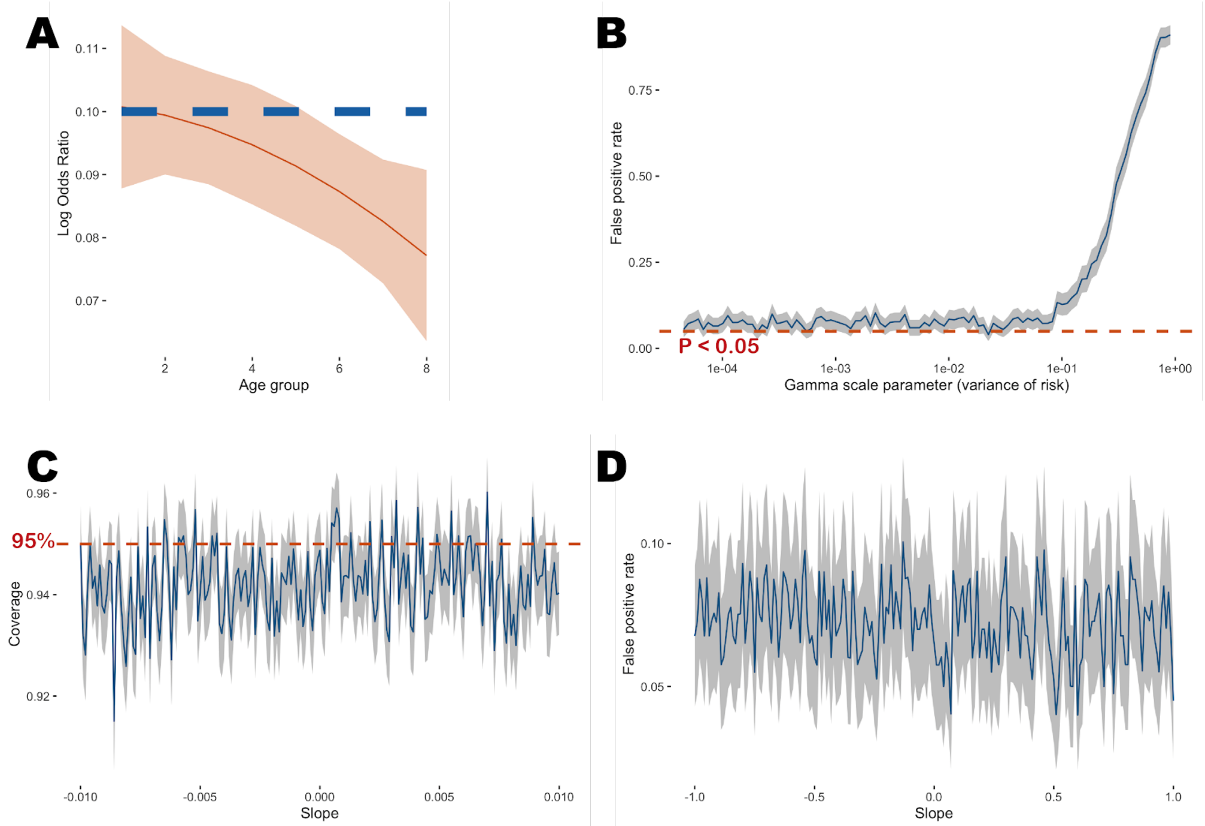


**Fig S2**: **Simulation studies.** A) A simulation with frailty showing that the inferred effect (red) deviates from the underlying effect size (blue dashed line). The variance of frailty in this case is 0.82. B) Effect of frailty on the false positive rate. The x-axis shows the variance of the frailty distribution, with a larger variance indicating stronger frailty, while the y-axis is the false positive rate of rejecting the true model of constant effect over age. The inferred curve does not deviate from uniformity when the frailty variance is smaller than 0.1. C) Coverage analysis. The blue curve shows the probability that 95% posterior credible interval covers the true genetic profile and the shaded area is the 95% confidence interval of the coverage estimate. D) Simulation to test the impact of selecting healthier individuals of older age. Selection bias towards healthier older people is simulated by changing the baseline hazard over age, such that a negative slope indicates a population in which older people are biased away from having disease. The blue solid line shows the false positive rate of rejecting the null hypothesis of uniformity for a baseline hazard with different slopes; the shaded area shows the 95% confidence interval. Genetic profile estimation uses a quadratic polynomial throughout.

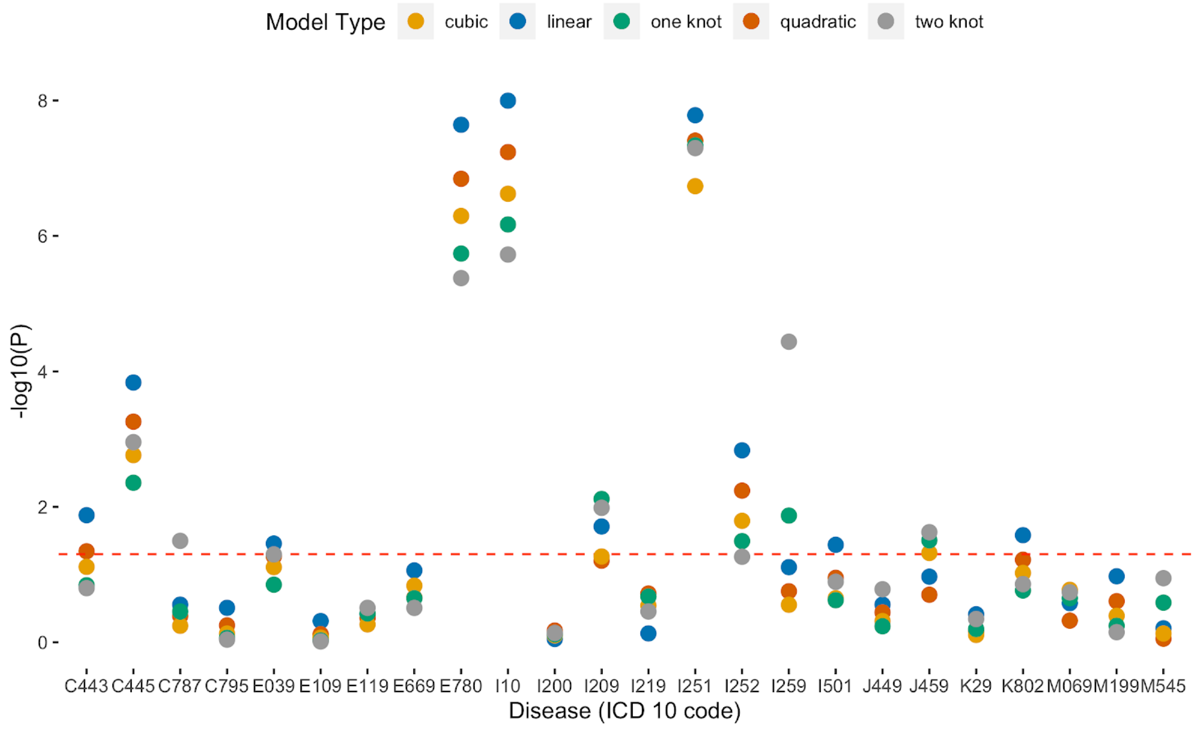


**Fig S3**: **Analysis of empirical data with curves of increasing complexity.** A likelihood ratio test is performed against a constant effect model (DF = 1) over age, for models with different smoothness. Smoothness is controlled by the degree of freedom of the spline basis, where we tested linear (DF = 2, blue), quadratic polynomial (DF = 3, red), cubic polynomial (DF = 4, orange), spline with one knot (DF = 5, green) and spline with two knots (DF = 6, grey). The red dotted line indicates P = 0.05.

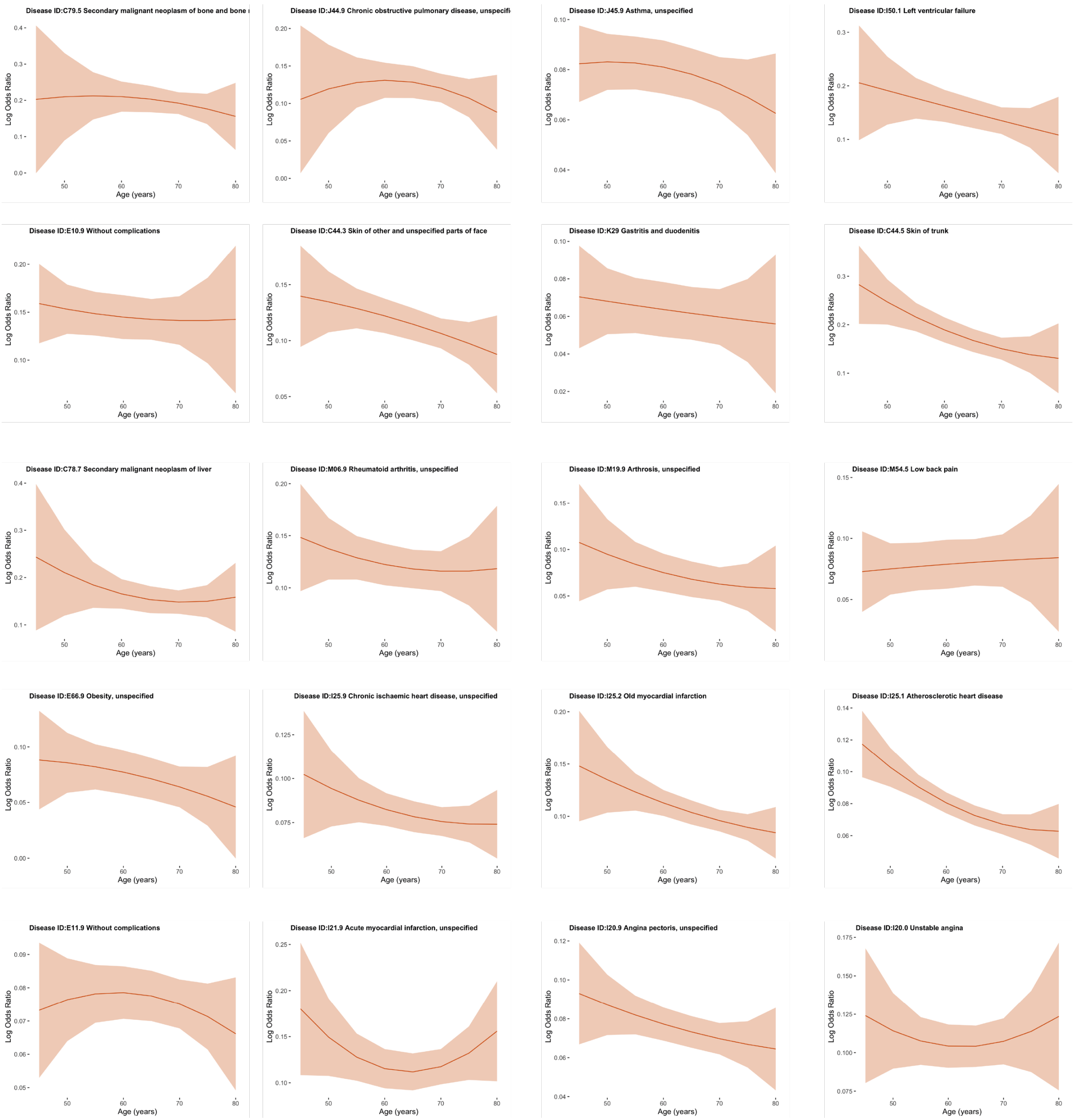


**Fig S4**: **Posterior estimation of genetic risk over age for all diseases**. The solid red curve indicates the posterior mean, and the shaded region is the 95% credible interval.

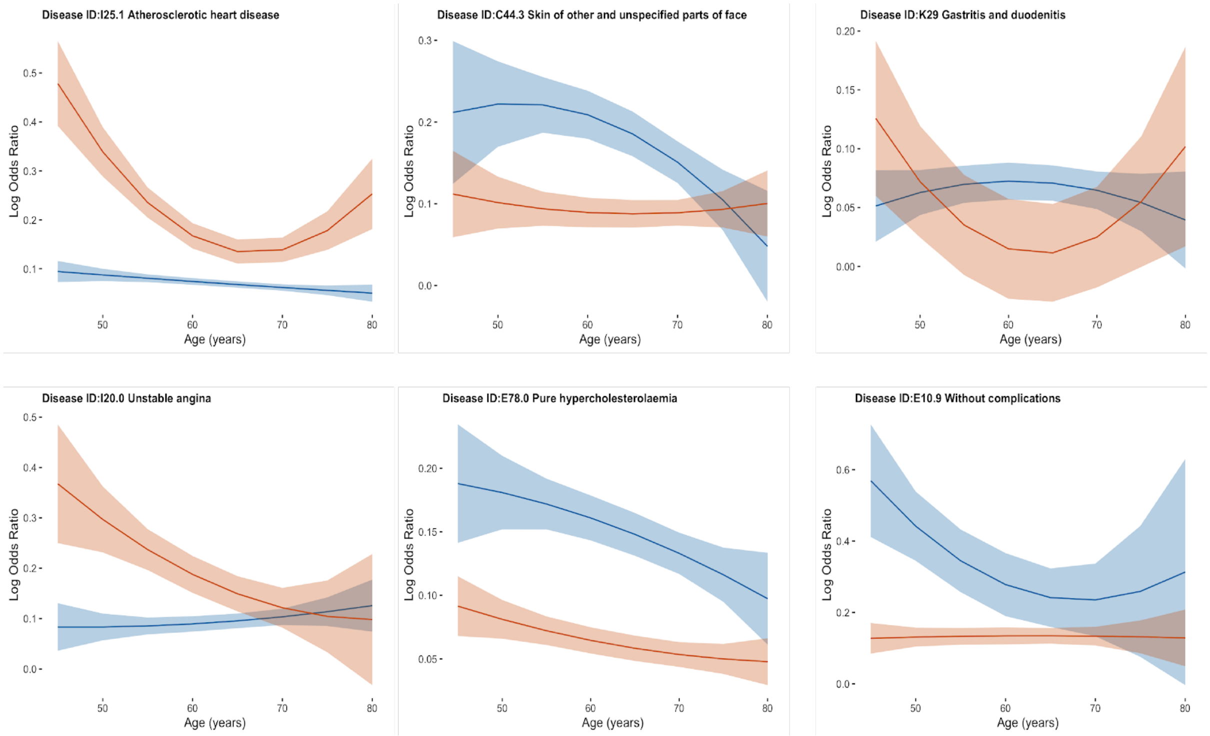


**Fig S5**: **Posterior curve estimates when two latent profiles are fitted for six diseases with moderate evidence of multiple profiles (P < 0.1).** The red and blue curves with corresponding shades show the profile means and 95% credible intervals for each profile.

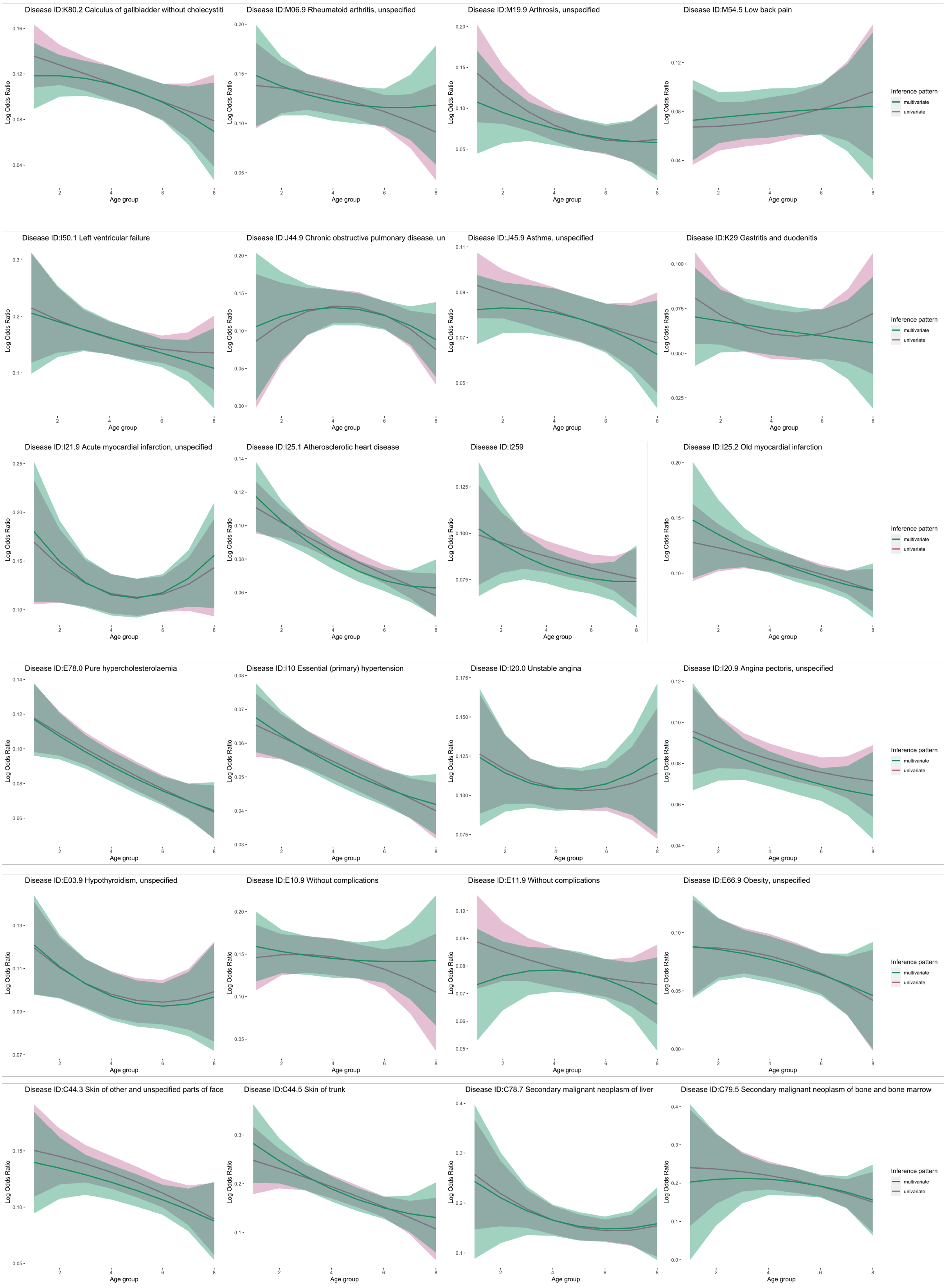


**Fig S6**: **Comparison of effect size estimation using multivariate and univariate methods for all diseases studied.** A quadratic polynomial model is fitted to the estimated effect size in both cases, which is shown as two curves: green (multivariate) and purple (univariate).

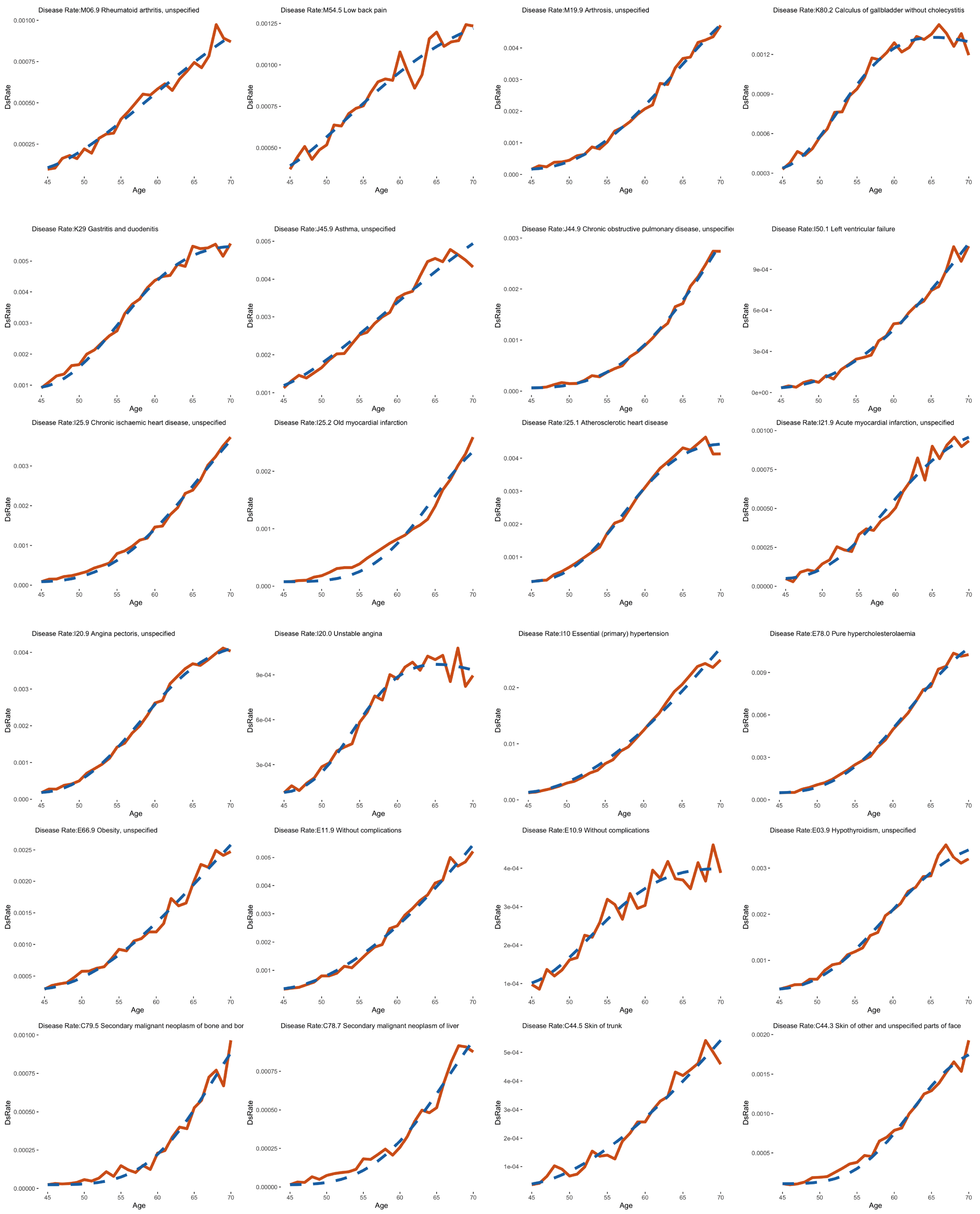


**Fig S7**: **Estimated incidence by age in the UK Biobank for all diseases studied here.** The red solid line indicates the rate estimated from the UK Biobank (see Methods) and the dotted blue line indicates the fitted incidence curve from the parametric model.

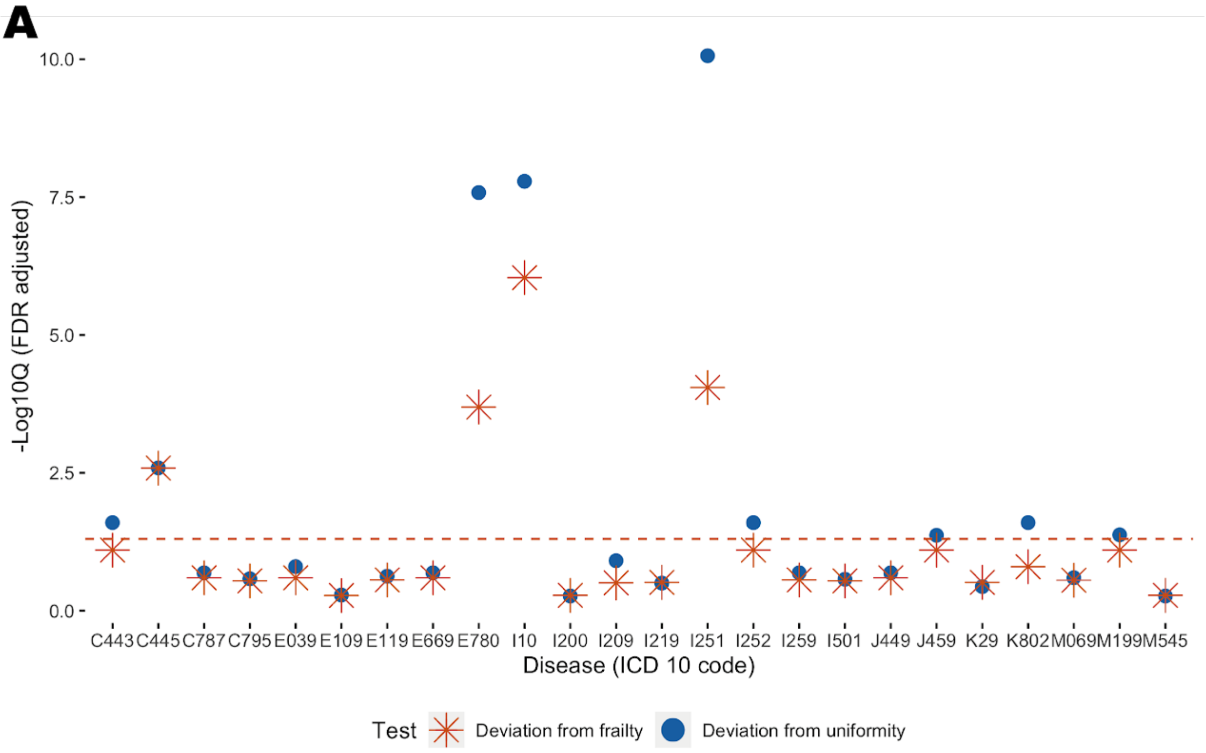


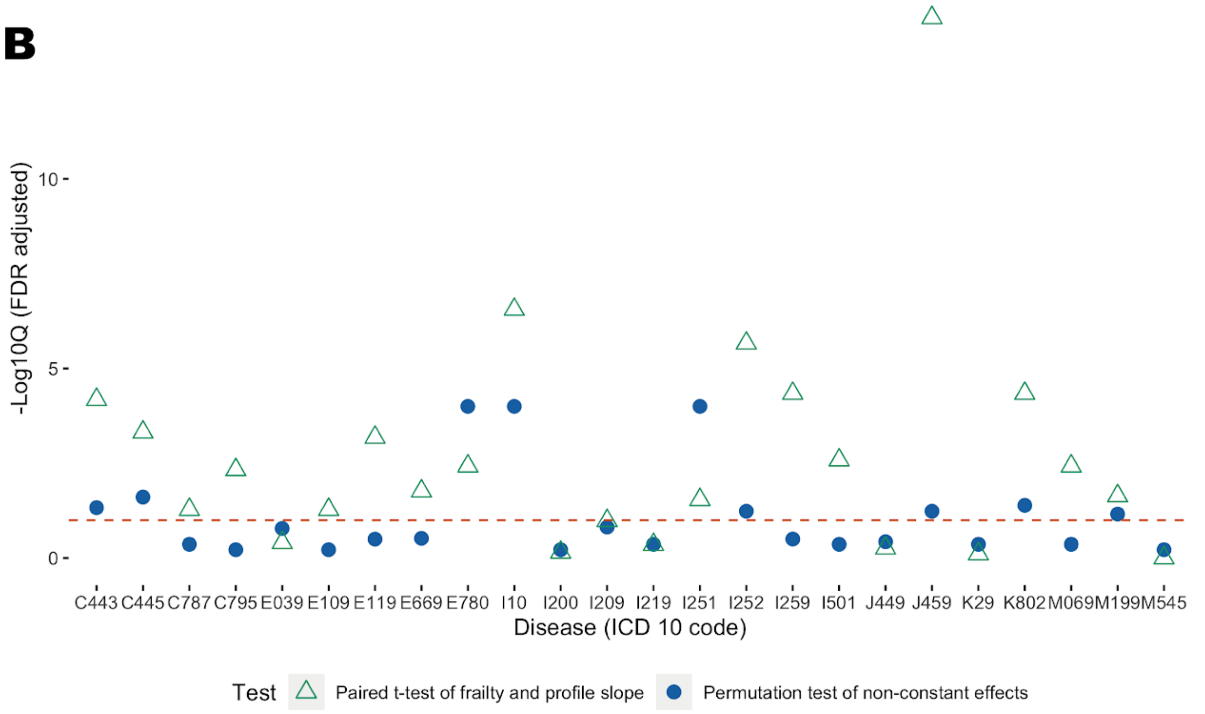


**Fig S8: Comparison of inferred genetic risk profiles and those predicted from fitted frailty models.** A) A likelihood ratio test of deviation from the fitted frailty model (red), compared with the likelihood ratio test of deviation from a constant effect model. Four diseases have Q < 0.05 after correcting for multiple testing. All inferences are performed on the univariate estimation of variant effect size because the fitted frailty should include both genetic and non-genetic factors. B) Paired t-test of the gradient of frailty and our inferred curve, identifying 17 out of the 24 diseases analysed where the inferred genetic risk profile slope is steeper than that implied by the inferred frailty parameters.

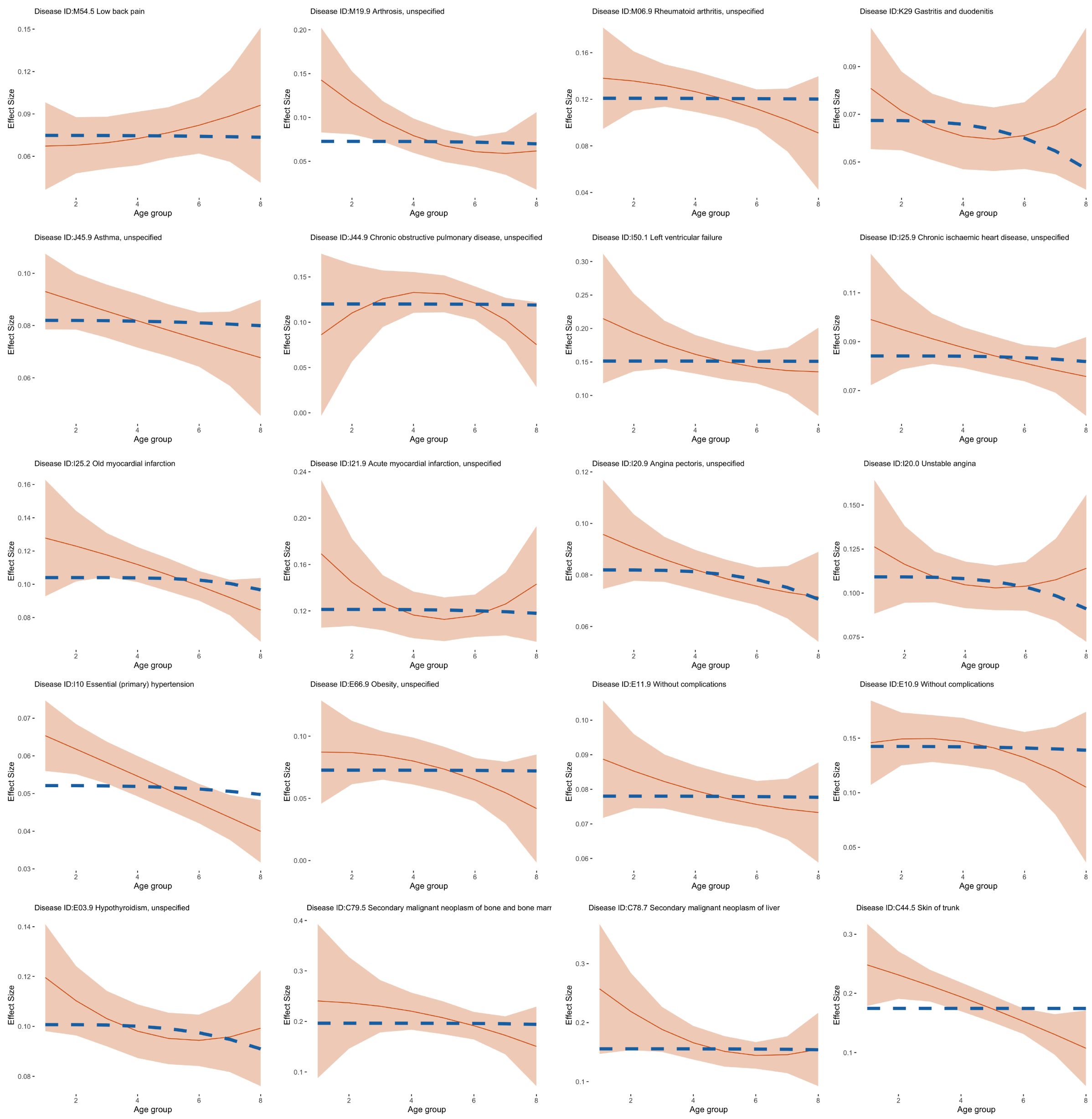


**Fig S9: Comparison of genetic risk estimated using quadratic polynomial and that predicted by a frailty model.** Comparison of fitted latent curves (red curve for the mean and shaded region for the 95% credible interval, estimated using the univariate approach) and latent profiles implied by the fitted frailty effect (blue dashed line), for all 24 diseases analysed here.

| ICD-10 code | Description | Prevalence in UK Biobank | Number of associated variants | Mean reported age of onset | P1 | Q1 | P2 | Q2 |
|---|---|---|---|---|---|---|---|---|
| C44.3 | Other and unspecified malignant neoplasm of skin of other and unspecified parts of face | 1.41% | 34 | 62.64 | 0.0118* | 0.0469* | 0.0024** | 0.0277* |
| C44.5 | Other and unspecified malignant neoplasm of skin of trunk | 0.48% | 25 | 61.45 | 0.0042** | 0.0247* | 0.6721 | 0.9003 |
| C78.7 | Secondary malignant neoplasm of liver and intrahepatic bile duct | 0.64% | 30 | 64.20 | 0.3367 | 0.4328 | 0.1558 | 0.5339 |
| C79.5 | Secondary malignant neoplasm of bone and bone marrow | 0.52% | 29 | 64.56 | 0.5859 | 0.6035 | 0.6863 | 0.9003 |
| E03.9 | Hypothyroidism, unspecified | 3.46% | 32 | 60.61 | 0.0760 | 0.1657 | 0.4978 | 0.7964 |
| E10.9 | Type 1 diabetes mellitus without complications | 0.63% | 28 | 57.32 | 0.5327 | 0.6035 | 0.2084 | 0.6250 |
| E11.9 | Type 2 diabetes mellitus without complications | 4.38% | 75 | 61.95 | 0.1853 | 0.3176 | 0.7878 | 0.9003 |
| E66.9 | Obesity, unspecified | 2.46% | 21 | 60.71 | 0.1514 | 0.3027 | 0.0372* | 0.1782 |
| E78.0 | Pure hypercholesterolemia | 8.12% | 43 | 62.59 | 0.0001** | 0.0001** | 0.0178* | 0.1063 |
| I10 | Essential (primary) hypertension | 18.98% | 80 | 61.87 | 0.0001** | 0.0001** | 0.3446 | 0.7685 |
| I20.0 | Unstable angina | 1.26% | 46 | 59.92 | 0.6035 | 0.6035 | 0.6899 | 0.9003 |
| I20.9 | Angina pectoris, unspecified | 3.84% | 78 | 61.84 | 0.0640 | 0.1535 | 0.8511 | 0.9046 |
| I21.9 | Acute myocardial infarction, unspecified | 0.92% | 24 | 62.08 | 0.3287 | 0.4328 | 0.3861 | 0.7721 |
| I25.1 | Atherosclerotic heart disease of native coronary artery | 4.46% | 116 | 61.57 | 0.0001** | 0.0001** | 0.0015** | 0.0277* |
| I25.2 | Old myocardial infarction | 1.72% | 83 | 64.32 | 0.0173* | 0.058* | 0.4493 | 0.7964 |
| I25.9 | Chronic ischaemic heart disease, unspecified | 2.66% | 86 | 63.69 | 0.1713 | 0.3162 | 0.2541 | 0.6774 |
| I50.1 | Left ventricular failure, unspecified | 0.83% | 22 | 63.22 | 0.3607 | 0.4328 | 0.9894 | 0.9894 |
| J44.9 | Chronic obstructive pulmonary disease, unspecified | 1.78% | 24 | 64.40 | 0.2318 | 0.3708 | 0.7321 | 0.9003 |
| J45.9 | Other and unspecified asthma | 6.34% | 35 | 57.54 | 0.0194* | 0.058* | 0.4901 | 0.7964 |
| K29 | Gastritis and duodenitis | 7.02% | 33 | 58.87 | 0.3452 | 0.4328 | 0.0167* | 0.1063 |
| K80.2 | Calculus of gallbladder without cholecystitis | 2.11% | 26 | 57.99 | 0.0086* | 0.0409* | 0.1287 | 0.5145 |
| M06.9 | Rheumatoid arthritis, unspecified | 0.99% | 54 | 60.16 | 0.3057 | 0.4328 | 0.7859 | 0.9003 |
| M19.9 | Osteoarthritis, unspecified site | 3.70% | 22 | 62.80 | 0.0261* | 0.0694* | 0.3523 | 0.7685 |
| M54.5 | Low back pain | 1.90% | 29 | 56.75 | 0.5659 | 0.6035 | 0.8669 | 0.9046 |

P1: permutation test for non constant profile over age; P2: permutation test for multiple profiles over age **P < 0.005 *P < 0.05
Q1: FDR adjusted P1 value; Q2: FDR adjusted P2 value **Q < 0.01 *Q < 0.1

**Table S1**: Summary of ICD-10 disease codes analysed and evidence for age-varying effect sizes and number of age-profile classes, fitted with a univariate model and quadratic polynomial. Details are for Table 1.

| ICD-10 code | Description | Starting risk size (intercept) | Profile Slope | Change per year | Genererti risk ratio of 75+ age group to 0-45 age group |
|---|---|---|---|---|---|
| C44.3 | Other and unspecified malignant neoplasm of skin of other and unspecified parts of face | 0.152 | -0.0076 | -1.00% | 63.18% |
| C44.5 | Other and unspecified malignant neoplasm of skin of trunk | 0.280 | -0.0215 | -1.53% | 41.83% |
| C78.7 | Secondary malignant neoplasm of liver and intrahepatic bile duct | 0.200 | -0.0079 | -0.79% | 71.03% |
| C79.5 | Secondary malignant neoplasm of bone and bone marrow | 0.248 | -0.0097 | -0.78% | 71.41% |
| E03.9 | Hypothyroidism, unspecified | 0.115 | -0.0038 | -0.65% | 76.37% |
| E10.9 | Type 1 diabetes mellitus without complications | 0.159 | -0.0031 | -0.39% | 86.16% |
| E11.9 | Type 2 diabetes mellitus without complications | 0.082 | -0.0012 | -0.29% | 89.75% |
| E66.9 | Obesity, unspecified | 0.099 | -0.0059 | -1.19% | 55.60% |
| E78.0 | Pure hypercholesterolemia | 0.120 | -0.0074 | -1.23% | 54.26% |
| I10 | Essential (primary) hypertension | 0.069 | -0.0036 | -1.06% | 61.01% |
| I20.0 | Unstable angina | 0.109 | -0.0004 | -0.08% | 97.15% |
| I20.9 | Angina pectoris, unspecified | 0.094 | -0.0040 | -0.85% | 68.91% |
| I21.9 | Acute myocardial infarction, unspecified | 0.129 | -0.0016 | -0.24% | 91.47% |
| I25.1 | Atherosclerotic heart disease of native coronary artery | 0.112 | -0.0075 | -1.33% | 50.06% |
| I25.2 | Old myocardial infarction | 0.146 | -0.0083 | -1.13% | 58.07% |
| I25.9 | Chronic ischaemic heart disease, unspecified | 0.096 | -0.0033 | -0.69% | 74.80% |
| I50.1 | Left ventricular failure, unspecified | 0.218 | -0.0139 | -1.27% | 52.40% |
| J44.9 | Chronic obstructive pulmonary disease, unspecified | 0.151 | -0.0056 | -0.74% | 73.18% |
| J45.9 | Other and unspecified asthma | 0.088 | -0.0022 | -0.50% | 82.00% |
| K29 | Gastritis and duodenitis | 0.072 | -0.0021 | -0.58% | 79.13% |
| K80.2 | Calculus of gallbladder without cholecystitis | 0.132 | -0.0060 | -0.91% | 66.57% |
| M06.9 | Rheumatoid arthritis, unspecified | 0.144 | -0.0047 | -0.65% | 76.33% |
| M19.9 | Osteoarthritis, unspecified site | 0.103 | -0.0067 | -1.29% | 51.87% |
| M54.5 | Low back pain | 0.072 | 0.0017 | 0.48% | 116.53% |

**Table S2**: Summary of changes in genetic risk contributions from before 45 years old to after 75 years old, when risk profiles are fitted using a linear model.

| ICD-10 code | < 45 years old | 45-50 years old | 50-55 years old | 55-60 years old | 60-65 years old | 65-70 years old | 70-75 years old | > 75 years old |
|---|---|---|---|---|---|---|---|---|
| C44.3 | 0.14 (SE = 0.023) | 0.135 (SE = 0.014) | 0.129 (SE = 0.009) | 0.122 (SE = 0.008) | 0.115 (SE = 0.007) | 0.106 (SE = 0.007) | 0.097 (SE = 0.009) | 0.088 (SE = 0.017) |
| C44.5 | 0.283 (SE = 0.04) | 0.247 (SE = 0.023) | 0.216 (SE = 0.015) | 0.189 (SE = 0.013) | 0.168 (SE = 0.012) | 0.151 (SE = 0.011) | 0.138 (SE = 0.019) | 0.131 (SE = 0.036) |
| C78.7 | 0.243 (SE = 0.077) | 0.21 (SE = 0.045) | 0.185 (SE = 0.024) | 0.166 (SE = 0.016) | 0.153 (SE = 0.014) | 0.148 (SE = 0.012) | 0.15 (SE = 0.017) | 0.158 (SE = 0.036) |
| C79.5 | 0.203 (SE = 0.102) | 0.21 (SE = 0.06) | 0.212 (SE = 0.033) | 0.21 (SE = 0.021) | 0.204 (SE = 0.018) | 0.192 (SE = 0.015) | 0.176 (SE = 0.021) | 0.156 (SE = 0.046) |
| E03.9 | 0.121 (SE = 0.012) | 0.111 (SE = 0.007) | 0.103 (SE = 0.006) | 0.097 (SE = 0.006) | 0.094 (SE = 0.005) | 0.093 (SE = 0.005) | 0.094 (SE = 0.007) | 0.097 (SE = 0.012) |
| E10.9 | 0.159 (SE = 0.021) | 0.153 (SE = 0.013) | 0.148 (SE = 0.011) | 0.145 (SE = 0.011) | 0.142 (SE = 0.011) | 0.141 (SE = 0.013) | 0.141 (SE = 0.022) | 0.142 (SE = 0.038) |
| E11.9 | 0.073 (SE = 0.01) | 0.076 (SE = 0.006) | 0.078 (SE = 0.004) | 0.079 (SE = 0.004) | 0.078 (SE = 0.004) | 0.075 (SE = 0.004) | 0.071 (SE = 0.005) | 0.066 (SE = 0.008) |
| E66.9 | 0.088 (SE = 0.022) | 0.086 (SE = 0.013) | 0.082 (SE = 0.01) | 0.077 (SE = 0.01) | 0.071 (SE = 0.009) | 0.064 (SE = 0.009) | 0.056 (SE = 0.013) | 0.046 (SE = 0.023) |
| E78.0 | 0.117 (SE = 0.01) | 0.107 (SE = 0.007) | 0.098 (SE = 0.005) | 0.09 (SE = 0.004) | 0.082 (SE = 0.004) | 0.076 (SE = 0.004) | 0.07 (SE = 0.005) | 0.064 (SE = 0.008) |
| I10 | 0.068 (SE = 0.005) | 0.062 (SE = 0.004) | 0.058 (SE = 0.003) | 0.054 (SE = 0.003) | 0.05 (SE = 0.003) | 0.047 (SE = 0.003) | 0.044 (SE = 0.003) | 0.042 (SE = 0.004) |
| I20.0 | 0.124 (SE = 0.022) | 0.114 (SE = 0.012) | 0.108 (SE = 0.008) | 0.104 (SE = 0.007) | 0.104 (SE = 0.007) | 0.107 (SE = 0.007) | 0.114 (SE = 0.013) | 0.124 (SE = 0.024) |
| I20.9 | 0.093 (SE = 0.013) | 0.087 (SE = 0.008) | 0.082 (SE = 0.005) | 0.077 (SE = 0.004) | 0.073 (SE = 0.004) | 0.07 (SE = 0.004) | 0.067 (SE = 0.006) | 0.064 (SE = 0.011) |
| I21.9 | 0.18 (SE = 0.036) | 0.149 (SE = 0.021) | 0.128 (SE = 0.013) | 0.115 (SE = 0.011) | 0.112 (SE = 0.01) | 0.117 (SE = 0.01) | 0.132 (SE = 0.014) | 0.156 (SE = 0.027) |
| I25.1 | 0.117 (SE = 0.01) | 0.103 (SE = 0.006) | 0.09 (SE = 0.004) | 0.08 (SE = 0.003) | 0.073 (SE = 0.003) | 0.067 (SE = 0.003) | 0.064 (SE = 0.005) | 0.063 (SE = 0.009) |
| I25.2 | 0.148 (SE = 0.026) | 0.135 (SE = 0.016) | 0.123 (SE = 0.009) | 0.113 (SE = 0.006) | 0.104 (SE = 0.006) | 0.096 (SE = 0.005) | 0.089 (SE = 0.006) | 0.084 (SE = 0.012) |
| I25.9 | 0.102 (SE = 0.018) | 0.094 (SE = 0.011) | 0.088 (SE = 0.006) | 0.082 (SE = 0.005) | 0.078 (SE = 0.004) | 0.076 (SE = 0.004) | 0.074 (SE = 0.005) | 0.074 (SE = 0.01) |
| I50.1 | 0.206 (SE = 0.053) | 0.191 (SE = 0.032) | 0.177 (SE = 0.019) | 0.163 (SE = 0.015) | 0.149 (SE = 0.014) | 0.135 (SE = 0.012) | 0.121 (SE = 0.018) | 0.108 (SE = 0.036) |
| J44.9 | 0.105 (SE = 0.049) | 0.119 (SE = 0.029) | 0.128 (SE = 0.017) | 0.131 (SE = 0.012) | 0.128 (SE = 0.011) | 0.121 (SE = 0.01) | 0.107 (SE = 0.013) | 0.088 (SE = 0.025) |
| J45.9 | 0.082 (SE = 0.008) | 0.083 (SE = 0.006) | 0.083 (SE = 0.005) | 0.081 (SE = 0.005) | 0.078 (SE = 0.005) | 0.074 (SE = 0.005) | 0.069 (SE = 0.008) | 0.063 (SE = 0.012) |
| K29 | 0.07 (SE = 0.014) | 0.068 (SE = 0.009) | 0.066 (SE = 0.007) | 0.064 (SE = 0.007) | 0.062 (SE = 0.007) | 0.06 (SE = 0.007) | 0.058 (SE = 0.011) | 0.056 (SE = 0.018) |
| K80.2 | 0.118 (SE = 0.014) | 0.118 (SE = 0.009) | 0.116 (SE = 0.008) | 0.111 (SE = 0.008) | 0.104 (SE = 0.007) | 0.095 (SE = 0.008) | 0.083 (SE = 0.013) | 0.069 (SE = 0.021) |
| M06.9 | 0.148 (SE = 0.026) | 0.137 (SE = 0.015) | 0.129 (SE = 0.01) | 0.122 (SE = 0.01) | 0.118 (SE = 0.009) | 0.116 (SE = 0.01) | 0.116 (SE = 0.016) | 0.118 (SE = 0.03) |
| M19.9 | 0.107 (SE = 0.032) | 0.095 (SE = 0.019) | 0.084 (SE = 0.012) | 0.075 (SE = 0.01) | 0.068 (SE = 0.01) | 0.063 (SE = 0.009) | 0.059 (SE = 0.013) | 0.058 (SE = 0.023) |
| M54.5 | 0.073 (SE = 0.016) | 0.075 (SE = 0.01) | 0.077 (SE = 0.01) | 0.079 (SE = 0.01) | 0.08 (SE = 0.009) | 0.082 (SE = 0.011) | 0.083 (SE = 0.018) | 0.084 (SE = 0.03) |

**Table S3**: Posterior mean risk profiles for all diseases analysed here, fitted with a single quadratic polynomial. Standard errors are also provided. Values are the mean effect size within the age interval for individual variants.

| ICD-10 code | Description | θ | $\gamma$(x10^(-6)) | k | Goodness-of-fit P-value |
|---|---|---|---|---|---|
| C44.3 | Other and unspecified malignant neoplasm of skin of other and unspecified parts of face | 0.94 | 0.04 | 3.58 | 0.76 |
| C44.5 | Other and unspecified malignant neoplasm of skin of trunk | 0.05 | 2.38 | 1.66 | 0.71 |
| C78.7 | Secondary malignant neoplasm of liver and intrahepatic bile duct | 0.34 | 0.05 | 3.11 | 0.65 |
| C79.5 | Secondary malignant neoplasm of bone and bone marrow | 0.38 | 0.01 | 3.79 | 0.71 |
| E03.9 | Hypothyroidism, unspecified | 0.74 | 4.20 | 2.24 | 0.69 |
| E10.9 | Type 1 diabetes mellitus without complications | 1.36 | 3.80 | 1.65 | 0.64 |
| E11.9 | Type 2 diabetes mellitus without complications | 0.04 | 16.75 | 1.77 | 1.00 |
| E66.9 | Obesity, unspecified | 0.17 | 6.83 | 1.83 | 0.99 |
| E78.0 | Pure hypercholesterolemia | 0.51 | 3.37 | 2.65 | 1.00 |
| I10 | Essential (primary) hypertension | 0.07 | 67.23 | 1.86 | 0.82 |
| I20.0 | Unstable angina | 1.66 | 1.93 | 2.40 | 0.96 |
| I20.9 | Angina pectoris, unspecified | 0.71 | 5.59 | 2.28 | 1.00 |
| I21.9 | Acute myocardial infarction, unspecified | 0.68 | 1.05 | 2.31 | 0.97 |
| I25.1 | Atherosclerotic heart disease of native coronary artery | 0.97 | 4.44 | 2.46 | 1.00 |
| I25.2 | Old myocardial infarction | 0.69 | 0.02 | 3.76 | 0.07 |
| I25.9 | Chronic ischaemic heart disease, unspecified | 0.27 | 1.24 | 2.54 | 1.00 |
| I50.1 | Left ventricular failure, unspecified | 0.10 | 1.47 | 2.05 | 1.00 |
| J44.9 | Chronic obstructive pulmonary disease, unspecified | 0.14 | 0.43 | 2.75 | 1.00 |
| J45.9 | Other and unspecified asthma | 0.25 | 60.12 | 1.33 | 0.01 |
| K29 | Gastritis and duodenitis | 1.13 | 19.46 | 1.99 | 0.99 |
| K80.2 | Calculus of gallbladder without cholecystitis | 1.87 | 8.56 | 1.90 | 0.93 |
| M06.9 | Rheumatoid arthritis, unspecified | 0.24 | 9.45 | 1.44 | 0.99 |
| M19.9 | Osteoarthritis, unspecified site | 0.29 | 4.67 | 2.22 | 1.00 |
| M54.5 | Low back pain | 0.62 | 21.07 | 1.25 | 0.81 |

**Table S4**. Estimated parameters of frailty from the UK Biobank. The fitted model has a hazard rate of $h_i = u_i \gamma t^k$, where $u_i$ ~ Gamma(shape=1/θ, scale=θ).

| ICD-10 code | Description | Baseline incidence rate (age 45-50) | Baseline incidence rate (age 70-75) | Average genetic risk multiplier within top 10% genetic risk group (age 45-50) | Average genetic risk factor within top 10% genetic risk group (age 70-75) | Incidence rate within top 10% genetic risk group (age45-50) | Incidence rate within top 10% genetic risk group (age 70-75) |
|---|---|---|---|---|---|---|---|
| C44.3 | Other and unspecified malignant neoplasm of skin of other and unspecified parts of face | 0.06% | 0.62% | 3.12 | 1.81 | 0.20% | 1.11% |
| C44.5 | Other and unspecified malignant neoplasm of skin of trunk | 0.03% | 0.16% | 6.34 | 3.88 | 0.21% | 0.61% |
| C78.7 | Secondary malignant neoplasm of liver and intrahepatic bile duct | 0.02% | 0.36% | 9.88 | 1.66 | 0.19% | 0.60% |
| C79.5 | Secondary malignant neoplasm of bone and bone marrow | 0.02% | 0.32% | 5.41 | 2.51 | 0.08% | 0.81% |
| E03.9 | Hypothyroidism, unspecified | 0.23% | 1.07% | 5.02 | 2.42 | 1.17% | 2.60% |
| E10.9 | Type 1 diabetes mellitus without complications | 0.06% | 0.12% | 4.42 | 2.20 | 0.25% | 0.26% |
| E11.9 | Type 2 diabetes mellitus without complications | 0.22% | 1.66% | 2.17 | 1.60 | 0.47% | 2.66% |
| E66.9 | Obesity, unspecified | 0.19% | 0.86% | 1.77 | 1.32 | 0.33% | 1.13% |
| E78.0 | Pure hypercholesterolemia | 0.31% | 3.42% | 2.59 | 1.64 | 0.81% | 5.60% |
| I10 | Essential (primary) hypertension | 0.88% | 8.02% | 2.14 | 1.95 | 1.88% | 15.66% |
| I20.0 | Unstable angina | 0.08% | 0.27% | 16.04 | 2.04 | 1.26% | 0.55% |
| I20.9 | Angina pectoris, unspecified | 0.15% | 1.29% | 10.28 | 1.69 | 1.56% | 2.17% |
| I21.9 | Acute myocardial infarction, unspecified | 0.04% | 0.35% | 3.10 | 1.73 | 0.11% | 0.61% |
| I25.1 | Atherosclerotic heart disease of native coronary artery | 0.18% | 1.38% | 6.62 | 2.40 | 1.22% | 3.31% |
| I25.2 | Old myocardial infarction | 0.05% | 1.05% | 6.55 | 1.92 | 0.34% | 2.03% |
| I25.9 | Chronic ischaemic heart disease, unspecified | 0.08% | 1.35% | 8.24 | 1.62 | 0.70% | 2.19% |
| I50.1 | Left ventricular failure, unspecified | 0.03% | 0.40% | 3.01 | 1.81 | 0.08% | 0.72% |
| J44.9 | Chronic obstructive pulmonary disease, unspecified | 0.05% | 0.95% | 2.84 | 1.42 | 0.14% | 1.34% |
| J45.9 | Other and unspecified asthma | 0.68% | 1.34% | 2.31 | 1.75 | 1.57% | 2.34% |
| K29 | Gastritis and duodenitis | 0.63% | 1.76% | 1.32 | 1.06 | 0.82% | 1.87% |
| K80.2 | Calculus of gallbladder without cholecystitis | 0.21% | 0.41% | 1.46 | 1.49 | 0.30% | 0.61% |
| M06.9 | Rheumatoid arthritis, unspecified | 0.07% | 0.30% | 4.92 | 2.43 | 0.35% | 0.73% |
| M19.9 | Osteoarthritis, unspecified site | 0.14% | 1.60% | 1.76 | 1.24 | 0.25% | 1.98% |
| M54.5 | Low back pain | 0.22% | 0.38% | 1.73 | 1.46 | 0.39% | 0.55% |

**Table S5**. Comparison of baseline hazard rate (population-level risk), genetic risk factor effect size and absolute hazard rate (baseline hazard multiplied by genetic risk factor) for an early age group and a late age group across the diseases studied here. The genetic risk factor and absolute hazard are computed from the group with the highest decile of genetic risk.
