## Supplemental Methods for "The impact of age on genetic risk for common diseases"

December 16, 2020

#### Contents

**1 Estimating genetic risk effect size over age**
1.1 UK Biobank dataset . . . . . . . . . . . . . . . . . . . . . . . . . . . . .
1.2 Interval censored proportional hazard model for genetic effect estimation . .
1.2.1 Constructing interval censored data sets . . . . . . . . . . . . . . . . .
1.2.2 Selecting variants of interest . . . . . . . . . . . . . . . . . . . . . . .
1.2.3 Effect size estimation . . . . . . . . . . . . . . . . . . . . . . . . . . .

**2 Identification of genetic effect heterogeneity over age**
2.1 Bayesian clustering of genetic risk profiles . . . . . . . . . . . . . . . . . .
2.1.1 Inference of genetic risk over age . . . . . . . . . . . . . . . . . . . .
2.2 Estimating effects of unobserved risk background . . . . . . . . . . . . . . .
2.2.1 Estimating unobserved risk background from incidence rates . . . .
2.2.2 Predicting the impact of frailty on age-dependent genetic risk . . . .
2.3 Statistical tests . . . . . . . . . . . . . . . . . . . . . . . . . . . . . . . .
2.3.1 Choice of latent curve degree of freedom . . . . . . . . . . . . . . . .
2.3.2 Permutation testing for genetic effect heterogeneity over age . . . . .
2.3.3 Testing for multiple clusters of variants . . . . . . . . . . . . . . . .
2.3.4 Test against a general frailty model . . . . . . . . . . . . . . . . . . .
2.4 Estimating age-dependency of genetic risk score in prediction . . . . . . . .

**3 Simulation**
3.1 Simulating a disease data set from an underlying genetic risk profile . . . .
3.1.1 Identifying age-varying genetic effects . . . . . . . . . . . . . . . . .
3.1.2 Detection of multiple clusters . . . . . . . . . . . . . . . . . . . . . .
3.1.3 Frailty . . . . . . . . . . . . . . . . . . . . . . . . . . . . . . . . . . .
3.1.4 Coverage analysis . . . . . . . . . . . . . . . . . . . . . . . . . . . . .
3.1.5 Selection of healthier elder population . . . . . . . . . . . . . . . . .

3.2 Potential models for the decreasing risk effect with age . . . . . . . . . . . .
3.2.1 GxE and GxG interaction . . . . . . . . . . . . . . . . . . . . . . . .
3.2.2 A threshold model for disease . . . . . . . . . . . . . . . . . . . . . .

### 1 Estimating genetic risk effect size over age

In order to estimate genetic risk profiles over age, we designed a framework which consists of two parts. The first part captures the risk size over age of each genetic variant and the second part identifies the pattern of these profiles using a clustering of curves method.

Section 1.1 covers the data source and processing procedure. Section 1.2.3 describes our "interval-censored" proportional hazard model to estimate genetic risk coefficients for variants across a sequence of age intervals. Section 2.1 describes a clustering method to infer the genetic risk age profile for a single disease, including the generative model, EM-inference, posterior estimation. Section 2.3 describes the statistical testing procedures to identify genetic heterogeneity over age, including permutation tests and likelihood ratio tests. Section 3 describes the in silico Monte Carlo simulations performed to assess the power of our method and also to explore potential mechanisms underlying the observed age effects.

#### 1.1 UK Biobank dataset

Our analysis uses the genotype data, individual information and Hospital Episode Statistics (HES) data from 409,694 individuals of British Isles ancestry in the UK Biobank dataset. [1] 31 ICD-10 codes were identified with a prevalence $> 0.5\%$ and at least 20 independent associated variants as identified using the TreeWAS model [2]. Of these, we analysed 24 that correspond to specific disease conditions (as opposed to procedures) and that have sex- and age-distributions compatible with our framework. These are listed in Table 1.

For each ICD-10 code, we combine the primary and secondary diagnosis from the full HES data set. We use the starting date of the first episode that records the disease diagnosis to compute age for first disease onset, which is calculated as the difference between onset date to the month of birth (due to data privacy, we only have access to birth information specified to year and month). The onset age is then rounded to years. For each ICD-10 code, only the first recorded diagnosis of each individual is kept.

For the population under observation, we also computed the age at observation end-points, which is either the age of individuals at the last update of the data set (here 2018-02-14) or the age of death if a death event is recorded. We categorised the disease onset age into 8 age intervals, the first and last of which are "before 45 years old" and "after 75 years old", with 5-year intervals in between.

#### 1.2 Interval censored proportional hazard model for genetic effect estimation

##### 1.2.1 Constructing interval censored data sets

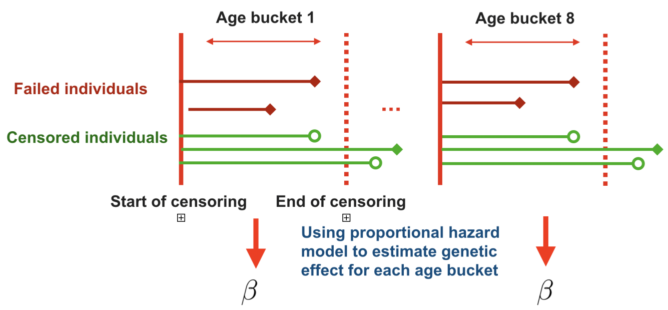


Figure 1: Schematic representation of an interval-censored data set. The red lines represent individuals who have disease onset within the age interval, while the green lines indicating individuals who have no onset within the interval. The rhombus represents a failure (disease event), while the circle represent a censoring event. Note that a disease event after the interval endpoint is considered as censored at the end of the interval.

For each disease of interest, we estimate the genetic effect size over 5-year age intervals (intervals are described in Section 1.1). Each age interval is an observation window of all healthy (alive and without onset of target disease) individual who survived pass the starting point of the interval. Onset of disease and exiting the study (death or no further records available) are recorded as "case" and "censored" events respectively. Events happening after this interval are considered right-censored at the end of the interval; see Figure 1.

We then perform case-control matching over the sub-population observed within each interval in two steps. First, we divide the sub-population into a disease group and control group. The disease group are those who have disease onset within the interval, and the control set are those who do not, including individuals who have disease onsets after the age interval so long as they remain healthy before the endpoint of the interval. This is what the term "interval-censored" means. In survival analysis, the control group are called "censored", and age at the "censoring" event is also needed for unbiased estimation. If a censoring event is observed within the age interval (i.e. the age of a death record or the last update in the UK Biobank is before the age interval end point) we use the age at the censoring event. If an individual does not have an event record within the age interval, we take their age at the end of the interval, regardless of their future events. Second, for each case in the disease group, we pick four nearest neighbors (without replacement) from the control group, matching sex, BMI, year of birth and 40 genetic principle components.

The covariates are available within the UK Biobank data set, over which we computed the principle components across the British Isle ancestry population. We then compute the Euclidean distance of the principle components to find the nearest neighbors in the population.

##### 1.2.2 Selecting variants of interest

The SNPs of interested are obtained through prior TreeWas analysis ($lg(BF_{tree}) > 5$, $BF_{tree}$ refer to Bayes factor of a single variant's effect over the model) [2]. We further filtered the set of SNPs to ensure LD-independent (loci kept with absolute Pearson correlation coefficient smaller than 0.2, $R^2 < 0.04$).

Our paper focuses on effect size changes over age. Therefore we used a standard GWAS approach to identify the risk and protective alleles at each locus [3], over the case-control matched dataset described in section 1.2.1. The first 40 genetic principle components are taken as covariates. For all loci that have protective minor allele (odds ratio ¡ 1), we switch allele labels to make sure we are estimating the effect size of risk alleles. Consequently, we make sure our clustering curve algorithms will discover variant clusters by different age-dependencies, not by protective or risk alleles.

##### 1.2.3 Effect size estimation

To obtain an unbiased estimate of genetic risk effect size over age, we use a proportional hazard (PH) model to estimate the genetic hazard ratio for different age groups, using the case-control matched data set described in Section 1.2.1.

Within each of the $M$ age intervals, we applied the Cox proportional hazard model to the disease group and control group, accounting the censoring effect. For the $m^{th}$ interval, a multivariate model is formulated as follows:

$$h(t_m|G) = h_0(t_m)exp\{\sum_{j=1}^{S} \hat{\beta}_j(t_m)G_j + \eta_m Z\}, \tag{1}$$

where $\hat{\beta}_j(t_m)$ is the target effect size of the $j^{th}$ variant in the $m^{th}$ interval and $\eta_m Z$ is the regression term of covariates. In our case $Z$ includes the first 40 genetics PCs of the UK Biobank and are regressed out for each interval. Both the mean and standard error of $\hat{\beta}_j(t_m)$ are obtained for each of the $M$ age intervals. These summary statistics are used subsequently for curve-cluster fitting.

To properly estimate the effect of frailty, we also use a univariate model to estimate the effect size of each $j^{th}(j = 1, 2, ..., S)$ SNP $G_j$ separately. This estimation includes the genetic background as part of frailty.

The estimation procedure described above is equivalent to a piece-wise proportional hazard model.

### 2 Identification of genetic effect heterogeneity over age

#### 2.1 Bayesian clustering of genetic risk profiles

In the sections before, we presented an unbiased framework for estimating risk over age for genetic variants. There are two hypotheses we want to test for common diseases. First, we want to test if there exist consistent non-constant genetic risk profiles. Second, we want to test if there are different clusters of genetic risk profiles. Therefore, we designed a Bayesian clustering of curves model, making use of the SNP-disease risk profiles obtained in Section 1.2.3.

##### 2.1.1 Inference of genetic risk over age

We applied a Bayesian clustering of curves model described in the Analytical Note. The model assumes each variant has a age-dependent effect profile which is generated from a mixture of curves model. The mean and standard error of the effect estimation procedure described above are the inputs of the model, from which we infer the underlying generative latent curve. The model allows vertical translation in the generative process (i.e. the likelihood won't change much if the profile of variant is far from the latent profile, as long as the shape of the variant curve is similar). The latent curve is a spline whose smoothness is controlled by changing the degrees of freedom. For detailed specification of model and hyper-parameters, see the Analytical Note.

Inference is performed by an EM algorithm (derived in the Analytical Note). Inference was repeated 20 times with random initialization of $\theta$ variable, where $\theta$ is sampled from a Gaussian distribution with mean 0 and standard deviation 0.0001. The highest likelihood sequence is retained. Since EM only provides a point estimate, we estimate the curve's credible interval using a variational approach. The derivation and proof of the approach are provided in Analytical Note.

#### 2.2 Estimating effects of unobserved risk background

##### 2.2.1 Estimating unobserved risk background from incidence rates

We assumed an individual hazard model that has a frailty coefficient $u$ and baseline hazard $h_0(t)$. The frailty coefficient has mean 1 and a scale parameter $\theta$ that controls the variance of population rate. We chose the baseline hazard to be a power function of age $t$.

$$\begin{aligned} &h(t|u) = u \cdot h_0(t) = u \cdot \gamma t^k \\ &u \sim \mathcal{G}(shape = \frac{1}{\theta}, scale = \theta) \; s.t. \; E[u] = 1, \; var[u] = \theta. \end{aligned} \tag{2}$$

We fit parameters of the above model from the population incidence, where the population hazard rate can be determined as (See Analytical Note):

$$h(t) = \frac{\gamma t^k}{1 + \theta\gamma \frac{t^{k+1}}{k+1}}. \tag{3}$$

We computed the empirical incidence rate $\hat{h}(t)$ in the UK Biobank. The empirical incidence rate at specific age is computed as the number of individuals who have first onset of the target disease within this age year, divided by the number of healthy individual at risk at the beginning of this age year. We then fit the parametric hazard in Equation 3 to the empirical incidence rate until age 70, and finally subtract the intercept from the empirical incidence rate to match the parametric form of hazard rate. We fit to minimize square error using the Nelder-Mead method,

$$argmin_{k,b,\gamma} \sum_t (\hat{h}(t) - h(t))^2$$

. The fitted incidence curves are compared with empirical curves for all diseases (Figure S7). We also computed a Goodness-of-fit p-value for each disease, comparing the match between fitted and empirical 3-year incidence rates using a Chi-square test statsitic. The Goodness-of-fit p-values are shown in Table S4.

##### 2.2.2 Predicting the impact of frailty on age-dependent genetic risk

Disease risk is influenced by many factors including those that are not observed. The unobserved confounding variables can create biases for estimators of effect size [4]. We show the how unmeasured variables can affect genetic risk effect size estimation, even when independent of the genetic risk factors (see Analytical Note).

Using the same model for frailty as in Section 2.2, we formulate the proportional hazard model as follows:

$$h(t|x, u) = h_0(t)e^{\beta x}u,$$

Where $h(t|x, u)$ is the conditional hazard rate, $x$ is the focal genetic variant of interest and $u$ is the unobserved risk (frailty). Instead of estimating the effect size $\beta$ and baseline risk $h_0(t)$ under the true model specification, we could only estimate marginal effect size $\beta^*$ and baseline hazard $h_0^*(t)$ under a misspecified model:

$$h(t|x) = h_0^*(t)e^{\beta^* x}.$$

Using the same parametric form of $u$ as in Equation 2, we obtain the marginal distribution for $\beta^*$ and $h_0^*(t)$ directly. Since disease typically has a low incidence rate in the population, we assume $\Lambda(t) = \int_0^t h_0(t)dt$ and $\beta$ to be a small, which leads to an approximation for $\beta^*$:

$$\beta^* \approx \beta(1 - \theta \cdot \Lambda(t)). \tag{4}$$

The frailty parameters estimated from the incidence profiles (Table S4) are applied to Equation 4 to obtain a correction factor.

#### 2.3 Statistical tests

##### 2.3.1 Choice of latent curve degree of freedom

Our model fits latent curves using splines, the degree of freedom (DF) of which will control smoothness. In order to choose the best DF, we fit the model with different spline bases using the method described in Analytical Note and compute the maximum likelihood. Since the spline $X\theta_i$ with degrees of freedom $p$ is a linear combination of columns of basis $X(\in \mathbb{R}^{M\times p})$, it is strictly nested. The model with fewer degrees of freedom is equivalent to setting elements of $\theta$ to be zero. Therefore, we opt for a likelihood ratio test to choose the best degree of freedom. We compute the likelihood ratio of models with $DF = 2, 3, 4, 5, 6$ degrees of freedom compared to a constant effect model ($DF = 1$), The negative log p-value (assuming a Chi-squared distribution of test statistic) are shown in Figure S3. The models we consider are linear (DF = 2), quadratic polynomial (DF = 3, cubic polynomial (DF = 4), cubic spline with one knot (DF = 5), cubic spline with two knots (DF = 6), natural cubic spline model with one knots (DF = 3), natural cubic spline model with two knots (DF = 4).

The seven likelihood ratio tests are performed for each disease, and then we extract the rank of fitting for different tests by ordering the p values for different models. Linear model (DF = 2) and quadratic polynomial model (DF=3) have the best overall rank and outperform the other models by a substantial margin. Although the linear model has a higher mean rank, we also consider the quadratic polynomial (DF=3) to detect non-linear dependency of genetic risk size and age.

##### 2.3.2 Permutation testing for genetic effect heterogeneity over age

To provide robustness in testing for age heterogeneity, we carried out a permutation test, using the likelihood ratio test statistic (for fitting a non-uniform genetic risk over age) for both the original data set and permutation samples to obtain the permutation p-value.

The likelihood ratio test compares an alternative model with linear and quadratic genetic risk over age and a null model assuming a constant effect over age. To compute the likelihood of the null model, we fit a constant effect model by setting following derivative to zero:

$$\frac{\partial}{\partial \beta_0} \sum_{j=2}^{S} z_{1j} \ln \mathcal{N}(\hat{\beta}_j | \beta_0 \mathbf{1}, \Sigma_j) = 0$$

$$\beta_0 = \frac{\sum_{j=1}^{S} \mathbf{1}^T z_{1j} \Sigma_j^{-1} \hat{\beta}_j}{\sum_{j=1}^{S} \mathbf{1}^T z_{1j} \Sigma_j^{-1} \mathbf{1}}. \tag{5}$$

Here $\mathbf{1}(\in \mathbf{R}^M)$ is a constant vector of length $M$. Plugging $\beta_0\mathbf{1}$ into the likelihood function (see Analytical Note), we compute the likelihood of the constant genetic effect model. The alternative models are fitted with linear or quadratic polynomial using EM (see Analytical Note). We then compute the likelihood ratio of alternative model and null model.

To perform permutation tests, we kept the case-control structure in Section 1.2.1 and then sample case-control pairs for each age interval, while fixing the onset age distribution for permutation samples. We repeat the procedure 10,000 times to obtain permutation samples, and compute the likelihood ratio for each sample.

We note that the likelihood ratio does not include the prior term $p(\theta_i)$ for spline coefficients, while EM finds the Maximum a Posteriori (MAP) estimate (see Analytical Note), which will give a likelihood slightly lower than the MLE estimate. Under the permutation test framework, the p-values will be consistent as long as we use the same test statistics for both the original data set and permutation samples. [5] We further checked that the difference between MAP estimation and MLE estimation is negligible.

The EM procedure (see Analytical Note) is initialised randomly as described in Section 2.1.1. We correct for multiple testing using FDR, with the corrected q-values shown in Table 1 (when a multivariate approach is used to estimate effect size) and Table S1 (when a univariate approach is used). The inferred profiles with 95% credible interval are shown in Figure S4.

###### 2.3.3 Testing for multiple clusters of variants

In order to determine the optimal number of clusters for each disease, we perform permutation test using same procedure as Section 2.3.2, considering the addition of each new cluster. For adding the $(k+1)^{th}$ cluster, the alternative model has $(k+1)$ clusters and null model has $k$ clusters. All models are fitted with quadratic polynomials (see Analytical Note). Again, we computed the likelihood ratio statistics for both the observed data set and permutation samples to obtain p-values. This analysis is performed over all diseases and adjusted for multiple testing with FDR. We note that we found no compelling evidence supporting a model of more than two clusters for any disease. The p-values and q-values for the test of two clusters are shown in Table 1 and Table S1.

The inferred latent curves are plotted for several diseases in Figure S5, where the p-values for rejecting a single cluster model are smaller than 0.1.

###### 2.3.4 Test against a general frailty model

In section 2.2, we discussed the effects of unobserved risk factors (frailty) on effect size estimation and provided a way to estimate a parametric model of frailty using the empirical incidence curve. The frailty effect is estimated as an frailty multiplier $\alpha$ in Equation 4. We then compute the frailty multiplier $\alpha(\in \mathbb{R}^M) = (1 - \theta\Lambda(t))$ for each disease, which allows us to fit effect size $\beta_0$ with a frailty correction, similar to Equation 5:

$$\frac{\partial}{\partial\beta_0}\sum_{j=2}^{S} z_{1j} \ln \mathcal{N}(\hat{\beta}_j | \beta_0\alpha, \Sigma_j) = 0$$

$$\beta_0 = \frac{\sum_{j=1}^{S} z_{1j}\alpha^T \Sigma_j^{-1}\hat{\beta}_j}{\sum_{j=1}^{S} z_{1j}\alpha^T \Sigma_j^{-1}\alpha}.$$

Here, $\hat{\beta}_j$ and $\Sigma_j$ are the summary statistics inferred for each variant over age. Since a single genetic effect size is small compared to the overall risk, all single variants could be considered to share the same risk background, which includes the genetic background. Therefore, we perform univariate estimation for each variant, using the method described in Section 1.2.3. To be specific, instead of estimating the effect size for each interval with all variants, we estimate the effect of a single variant's effect each time:

$$h(t_m|G_j) = h_0(t_m)exp\{\hat{\beta}_j(t_m)G_j + \eta_m Z\},$$

where $Z$ is the first 40 genetic PCs, included to account for the population structure as described in Section 1.2.3.

To test if the inferred genetic risk profile deviates from the model predicted by the fitted frailty parameters, we infer the latent age profile accounting for frailty by multiplying $\alpha$ by the intercept base, i.e. in the model without frailty the first spline base is the intercept $X_1 = \mathbf{1}(\in \mathbf{R}^M)$, while the first base is the frailty multiplier $X_1 = \alpha$ in a model with frailty correction (see Analytical Note). We then compute the likelihood ratio test as in Section 3.1.1. In the frailty-corrected model the null model is fitted with a latent curve whose shape is solely determined by the frailty term $\alpha$, while the Alternative model is a quadratic polynomial with the intercept $X_1 = \alpha$. The test will show if the risk profile deviates from that predicted by frailty. After correcting for multiple testing using FDR, we plot q-values for the deviation from the frailty model, along with q-values for deviation from uniformity (also computed from likelihood ratio test) in Figure S8A.

Since the genetic risk shows a consistent decreasing pattern, we next sought to test if the decreasing pattern could be explained only by the frailty correction term. For each disease, we compute the gradient of the inferred latent curve and the fitted frailty curve (Figure S9), and performed a one-tailed paired t-test on whether the gradient of fitted curve is greater than the gradient caused by frailty. (Figure S8B) The results are corrected for multiple testing by FDR.

It is worth noting that estimation of the unobserved background effects (frailty) uses the population incidence rate over age, which is independent of our estimation of latent profiles.

#### 2.4 Estimating age-dependency of genetic risk score in prediction

To assess whether the collective effect of risk variants, as captured by a combined genetic risk score (or polygenic risk score - PRS), show similar profiles of age-varying risk, we used the case-control matching procedure described in section 1.2.1 with five-fold cross-validation, keeping 20% of case/controls for each age interval as test sets, and estimating effect sizes for the selected variants in the remaining 80% of case/controls using multivariate logistic regression (including age, sex and 40 genetic PCs). The effect of the PRS on risk within each age interval in the test set was then estimated (again with logistic regression and covariates). We estimated the odds-ratio for the top decile of risk and the top 20% of risk, using 20 repeats of the procedure to obtain the bootstrap sampling distribution.

### 3 Simulation

#### 3.1 Simulating a disease data set from an underlying genetic risk profile

In our simulation, we generate a risk age profile for each variant from underlying curves with different slopes. The individual risk is then computed at different ages, which are then used to generate disease incidence events over the simulated population.

We choose the population size to be 50,000, which is comparable to our empirical case-matching population size (the set of common diseases analysed each have 10,000 cases in the UK Biobank and we match each case with four controls). We simulated 50 SNPs (MAF of each SNPs are sampled from uniform distribution 0-0.5). The risk effect for each SNP is sampled from a profile which changes with age linearly. The individual hazard at a specific age interval $t_m$ is computed as the exponential of genetic risk multiplied by a linearly increasing baseline hazard ratio $h_0(t_m)$. The hazard for each individual within an interval is assumed to be constant.

$$h(t_m|G) = h_0(t_m) * exp\{\sum_{j=1}^{S} \beta_j(t_m) G_j\}. \tag{6}$$

For each interval $t_m$, we simulated the time to the next event using a homogeneous Poisson process with rate $h(t_m|G)$. An individual with no event in this interval is considered as observed (censored). We record only the first event as the onset of the disease. The simulation is performed over a 40 year duration divided into eight 5-year intervals, as most of the disease onset occurs between ages 40-80 years old in the UK Biobank. In order to represent the end of observation (study drop-out) or death events in the cohort, a competing censoring process is sampled using a Poisson process of constant rate $h_c$. The dropout/death and disease onset events are combined and we keep the first event, labelling it as either disease or censoring.

The parameters required for the simulation include: the censoring rate (per year) $h_c$, the number of intervals $M$, the number of individuals $N$, the number of variants $S$, the baseline hazard rate $h_0(t_m)$, the baseline genetic risk effect size $\beta_0$, the latent curve $\beta$ and the curve variance term $\Sigma$. We set these parameter for the simulation as follows:

$$\begin{aligned}
&h_c = 0.01 \\
&M = 8 \\
&N = 50000 \\
&S = 50 \\
&h_0(t_m) = (5 + t_m) * 10^{-4} \\
&\beta_0 = 0.1 \\
&\beta = \beta_0 + s * [-3.5, -2.5, -1.5, -0.5, 0.5, 1.5, 2.5, 3.5]^T \\
&\Sigma = 4 \times 10^{-4} + diag(10^{-4}) \\
&\hat{\beta}_j \sim \mathcal{N}(\beta, \Sigma)
\end{aligned} \tag{7}$$

We generate the $j^{th}$ variant's risk profile $\hat{\beta}_j$ from a normal distribution with mean $\beta$ and variance $\Sigma$. The variable $s$ controls the slope of the genetic risk profile, which varies between simulation studies.

###### 3.1.1 Identifying age-varying genetic effects

We simulated a population using the scheme described above and analysed it using the methods described in Sections 1.2.3 and 2.1 to infer the genetic risk profiles over age and the underlying curves that generate them. We simulated the cohort with different values of the slope, $s$, which represent different age dependencies, and tested whether our method could recover the simulated values. Examples of how fitted curves compare to the true curve are shown in Figure 3.

We then assessed the power of the statistical test described in Section 2.3.2. We simulated the genetic risk profile with the slope ranging from -0.01 (linearly decreasing with age) to 0.01 (linearly increasing with age), with a step size of $10^{-4}$. The simulated population is analysed using the null model of a constant effect with age, and an alternative model of either a linear model, or a quadratic polynomial curve. A likelihood ratio test is performed to calculate the p-value, and we calculate the power of rejecting the null at a threshold of $p \leq 0.05$. How the power changes with slope is shown in Figure 3. For each slope $s$, the simulation was repeated for 400 times to estimate the power and its standard error.

###### 3.1.2 Detection of multiple clusters

We use simulation to estimate the statistical power for detecting multiple clusters of genetic risk profiles. The disease cohort was simulated with parameter values as in Equation 7, except that five (10%) of the variants had effect sizes generated from a non-constant latent profile, while the effect sizes for the remaining 45 variants had a constant (age-invariant) effect of $\beta_0$. We assessed our model as to whether it can detect the presence of multiple clusters. The simulated cohorts are analysed with both a null model of a single quadratic polynomial curve, as well as the alternative model of two quadratic polynomial curves. Examples of fitted curves compared to the true curves are shown in Figure 3.

For each simulation, we compute the p-value for the likelihood ratio test comparing two clusters against one cluster, measuring power at $p \leq 0.05$. We varied the slope of the non-constant profile to test how different the curve needs to be from a constant effect to be distinguishable by our model. Power is computed for slope ranging between -0.0375 and 0.0375, with a step size of $2.5 \times 10^{-4}$. For each slope $s$, the simulation was repeated for 400 times to estimate the power and its standard error.

###### 3.1.3 Frailty

As described in Section 2.2, frailty can induce biases in the estimation of effect sizes with age and lead to rejection of the null hypothesis even if the risk are constant over age. We use simulation to estimate how the false positive rate changes with frailty.

To investigate this effect, we simulated the unobserved effect (frailty) $u$ as a multiplier to the disease hazard, and estimated the effect size under the mis-sepcified model (Equation 6) without $u$.

$$h(t_m|G) = h_0(t_m) * exp\{\sum_{j=1}^{S} \beta_j(t_m)G_j\} * u. \tag{8}$$

Here, each individual has its own frailty sampled from the gamma distribution with mean equal to one and scale parameter $\theta$ to specify the variance of the distribution:

$$u \sim Gamma(u|shape = \frac{1}{\theta}, scale = \theta).$$

We simulated the disease with a constant genetic risk effect over age and parameter settings as in Equation 7, with the individual hazard rate computed as in Equation 8. We then performed the estimation as in Section 3.1.1. The fitted curves diverge from the true underlying curve, as predicted by the analytical results in Section 2.2; an example is shown in Figure S2A with $\theta = 0.82$. The simulation was repeated for 400 times to estimate power (false positive rate) and its standard error.

###### 3.1.4 Coverage analysis

To estimate the posterior distribution of the latent curve, we adopt a Variational Bayesian approach to approximate the confidence interval (see Analytical Note). In order to evaluate the accuracy of the approximation, we performed a coverage analysis for the estimated confidence interval. Using the simulation results from Section 3.1.1, we computed coverage for the 95% credible interval (i.e. the proportion of true values that are inside the credible interval). The coverage rate is plotted against the slope of the genetic profiles in Figure S2C.

###### 3.1.5 Selection of healthier elder population

Previous research has showed that the UK Biobank has a selection bias towards healthier older people compared to the broader population [6]. In order to test whether this effect influences our results, we modelled such bias by modifying the baseline hazard function $h_0$ in Equation 6. Previously, we assumed the baseline hazard increases monotonically with age. To model a population with healthier older individuals we used a modified baseline:

$$h_0(t_m) = (9.5 + s * (t_m - 4.5)) * 10^{-4}.$$

Here $s$ is the slope of baseline hazard ranging between -1 and 1, with a step size of $10^{-2}$. We repeated the analyses described above and estimated the false positive rate as described in Section 3.1.1. The simulation was repeated 400 times at each value of $s$, and the result is shown in Figure S2D.

##### 3.2 Potential models for the decreasing risk effect with age

###### 3.2.1 GxE and GxG interaction

Across diseases, we observed a consistent pattern of a decreasing profile in the inferred genetic risk (Figure S4). To consider whether this pattern could be explained by interactions (either gene-by-environment or gene-by-gene) we performed additional simulations. We modelled the interaction of a focal genetic effect with other unobserved risk factors. Assuming the effect size $\beta$ interacts with environmental or other genetic factors, the effect size for each individual is generated from a positively defined distribution:

$$\beta \sim \mathcal{G}(\beta).$$

Following the proportional hazard model specified in Equation 1, we use $\beta^*$ to represent the marginal effect size under a mis-specified model:

$$h(t) = h_0(t) \exp(\beta^* x), \tag{9}$$

where $h_0(t)$ is a positive baseline hazard and $x$ is the covariate. We can show that the estimated marginal effect size $\beta^*$ will be increasingly underestimated as $t$ increases for all positive defined probability distributions for $\mathcal{G}(\beta)$ (see Analytical Note). We then performed a simulation using the parameter settings of Equation 7, but sampled an effect size $\beta$ for each individual from a gamma distribution $Gamma(shape = \frac{\beta_0}{\theta}, scale = \theta), \theta = 4$. The effect size for each individual remains a constant over age interval. We then inferred the posterior of effect size, presented in Figure 6B. We note that this model is a generalisation of the concept of frailty in which one allele has greater frailty than the other.

##### 3.2.2 A threshold model for disease

We simulated a threshold model in which each individual has an unknown "liability", which evolved over time [7]. For a specific disease, onset occurs when an individual's "liability" passes a certain threshold. We simulated a liability model for 50,000 individuals with a single genetic effect that alters the starting point of liability. Genotypes were simulated with a risk allele frequency 0.3.

The liability is simulated as a stochastic process with starting point at $L(t_0) = L_0 \times e^{-\beta_0 G}$, where $L_0 = -80$ is the population starting liability; $\beta_0 = 0.1$ is the effect size of genotype $G$. We then simulated $S$ increments of liability from a $S$ dimension Gaussian distribution $\mathcal{N}(\mathbf{1}^T, \Sigma_L)$, where $\Sigma_L = 4 \times (0.2 + diag[\mathbf{0.8}])$, controls the drift and variance of the stochastic process. The stochastic process models the disease risk increase over age through the drift $\mathbf{1}^T$, and the correlation of increments induced by $\Sigma_L$ creates a "momentum" such that an individual's health status tends to improve or deteriorate over years at similar rate. We simulated for $S = 60$ years and considered and individual to have an onset of a disease when the liability (arbitrarily) reaches 0. We then estimated the effect size of the risk allele over the age interval 21-60, using the methods described in Section 1.2.3.

One example of liability for each genotype is shown in Figure 6A, along with the estimated genetic effect size over age.
