## Supplementary material for "The impact of age on genetic risk for common diseases": Analytical Note

### Analytical Note for "Age modulates genetic risk for multiple common diseases"

October 9, 2020

#### Contents

**1 Bayesian clustering of curves**
1.1 Generative model specification . . . . . . . . . . . . . . . . . . . . . . . . .
1.2 Inference of cluster distribution: EM . . . . . . . . . . . . . . . . . . . . . . .
1.3 Approximating the confidence interval of latent curves: Variational inferences

**2 The effect of unobserved factors on risk effect estimation**
2.1 Independent unobserved effects: frailty . . . . . . . . . . . . . . . . . . . . . .
2.1.1 Parametric modelling of effect size in the presence of frailty . . . . .
2.1.2 Inferring frailty parameters from the population incidence rate . . .
2.2 Dependent unobserved effects: GxE and GxG interactions . . . . . . . . . . .

#### 1 Bayesian clustering of curves

##### 1.1 Generative model specification

We use a mixture of curves model to cluster genetic risk profiles to latent curves fitted with a spline basis. The graphic representation of the model is shown in Figure 1. Our analysis is performed independently for each disease, therefore without specific clarification, we omit the disease index.

Our model assumes risk profiles of the $S$ SNPs are assigned to $K$ clusters, where we denote the mean of risk profile for the $j^{th}$ SNP ($j = 1, 2, ..., S$) as $\hat{\beta}_j \in \mathbb{R}^M$ and the standard error of $\hat{\beta}_j$ to be $\epsilon_j \in \mathbb{R}^M$. $M$ refers to the number of age intervals we have. The mean and standard error of risk effect sizes within $M$ age intervals are the summary statistics which are our model inputs.

The model infers latent curves that generate risk profiles for each variant. We use a linear combination of cubic spline bases to construct smooth latent curves, where $X\theta_i(\in$

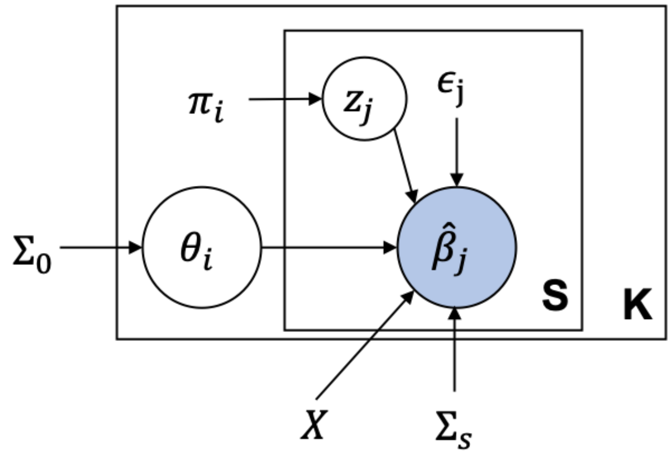


Figure 1: Schematic representation of the model. $i = 1, 2, .., K$ is the index of the mixture components; $j = 1, 2, .., S$ is the index of the SNPs. $\hat{\beta}_j$ and $\epsilon_j$ are the summary statistics for SNP $j$ that are estimated using an interval-censored proportional hazard model (See Methods). $z_j$ is the latent variable, which assigns SNP $j$ to cluster $i$.

$\mathbb{R}^M$) is the latent curve of $i^{th}$ cluster ($i = 1, 2, ..., K$). The cubic spline basis matrix $X \in \mathbb{R}^{M \times P}$ is defined as follows: [1, p. 141]

$$\begin{aligned} X_s(m) &= m^{s-1},\ s \leq 4 \\ X_s(m) &= (m - \xi_{s-4})^3_+,\ s > 4. \end{aligned} \tag{1}$$

Here $m = 1, 2, ..., M$, and $s = 1, 2, ...P$. $P$ are the number of bases, which controls the degrees of freedom of the cubic spline. $(\cdot)_+$ denotes the positive part of a function and $\xi$s are even split points across the $M$ age intervals. $\theta_i \in \mathbb{R}^P$ is the linear coefficients of $P$ bases.

Other variables shown in Figure 1 specify the generative process for the risk profiles, using a mixture model. $\Sigma_s$ is a vertical translation matrix with all elements equal, which allows $\hat{\beta}_j$ to be vertically translated from the latent curve $X\theta_i$ that generates it. $\pi_i$ is the mixture weight of the $i^{th}$ latent curve and $z_j \in \mathbb{R}^K$ is latent one-hot vectors assigning $j^{th}$ SNP to one of $K$ latent curves. We use $\Theta = (\pi, \epsilon, \theta, X, \Sigma_s)$ to denote the set of parameters in the mixture model, then we summarise our generative model as follows:

$$P(\hat{\beta}_j|\Theta) = \sum_{i=1}^{K} \pi_i \mathcal{N}(\hat{\beta}_j|X\theta_i, \Sigma_s + \Sigma_j). \tag{2}$$

Here $\Sigma_j \in \mathbb{R}^{M \times M}$ is a diagonal matrix with diagonal elements $\epsilon_j^2$. Intuitively, the model is a mixture of $K$ latent curves, each fitted with a spline $X\theta_i$. The profile for the $j^{th}$ SNP is generated from one of the latent curves with a variance $\Sigma_j$, then vertically translated with $\Sigma_s$. Note $\Sigma_j$ is the standard error for $\hat{\beta}_j$ (summary statistics; see Methods) and the vertical translation is controlled by a hyper-parameter $\Sigma_s$.

We further specify a prior distribution for $\theta_i$, which is a Gaussian distribution with zero mean and a fixed covariance matrix:

$$P(\theta_i) = \mathcal{N}(\theta_i|0, \Sigma_0). \tag{3}$$

In following sections we will derive the equations used in the inference of the latent curve $X\theta_i$ for each cluster. Note the spline bases $X$ are shared across clusters, so we infer the posterior of $\theta_i$.

Two hyper-parameters we have in the model are set as:

$$\Sigma_0 = diag(\mathbf{1}^P), \Sigma_s = 0.0004$$

#### 1.2 Inference of cluster distribution: EM

In our model, the variables for different clusters are exchangeable, [2] which will cause the so-called "label switching" problem if we apply a sampling method. Therefore, we derive an EM algorithm to maximize the following log likelihood:

$$l = \sum_{j=1}^{S} \ln \sum_{i=1}^{K} \pi_i \mathcal{N}(\hat{\beta}_j | X\theta_i, \Sigma_s + \Sigma_j) + \sum_{i=1}^{K} \ln \mathcal{N}(\theta_i|0, \Sigma_0). \tag{4}$$

To derive the inference equations we first write down the complete data log likelihood function with the latent variable $Z$:

$$\ln P(\hat{\beta}, \mathbf{Z}|\Theta) = \sum_{j=1}^{S} \sum_{i=1}^{K} z_{ji} \{\ln \pi_i + \ln \mathcal{N}(\hat{\beta}_j | X\theta_i, \Sigma_s + \Sigma_j)\}. \tag{5}$$

Classic mixture models use "cluster specific" covariance matrices, which are inferred for each mixture component. However, it is worth noting that we use "SNP specific" covariance matrices $\Sigma_j$ for the $j^{th}$ variant. The rationale is that when we estimate the effect size $\hat{\beta}_j$ for a specific locus, we obtain its standard error $\epsilon_j$. Therefore, we use different $\Sigma_j$ for each variant to capture the uncertainty differences across loci.

To keep the notation simple, from here on we will use $\Sigma_j$ to denote $\Sigma_s + \Sigma_j$. By maximizing the expected value of the complete data likelihood $E_z[\ln\{P(\hat{\beta}, Z|\Theta)P(\theta_i)\}]$ from Equation 5 and 3, we obtained the M-step update as follows:

$$\pi_i = \frac{N_i}{N},\ N_i = \sum_{j=1}^{s} \gamma(z_{ji}), \tag{6}$$

$$\theta_i = \{\Sigma_0^{-1} + \sum_{j=1}^{S} \gamma(z_{ji}) X^T \Sigma_j^{-1} X\}^{-1} \sum_{j=1}^{S} \gamma(z_{ji}) X^T \Sigma_j^{-1} \hat{\beta}_j. \tag{7}$$

$\gamma(z_{ji})$ is the expectation of $z_{ji}$, which will be obtained in the E-step. By Bayes' theorem, we could write down the posterior distribution of the latent variable $Z$:

$$P(\mathbf{Z}|\hat{\beta}, \Theta) \propto \prod_{j=1}^{S} \prod_{i=1}^{K} \{\pi_i \mathcal{N}(\hat{\beta}_j | X\theta_i, \Sigma_j)\}^{z_{ji}}. \tag{8}$$

Since the posterior factorizes over $j$, $\mathbf{z}_j$ follows an Multinoulli distribution for each SNP. From this we can derive the expectation of $z_{ji}$ , which is the E-step:

$$\gamma(z_{ji}) = E[z_{ji}] = \frac{\pi_i \mathcal{N}(\hat{\beta}_j | X\theta_i, \mathbf{z_j})}{\sum_{m=1}^{\mathcal{K}} \pi_m \mathcal{N}(\hat{\beta}_j | X\theta_m, \mathbf{z_m})}. \tag{9}$$

The inference is performed by randomly initialising $\theta_i$s, then alternating between the M-step and E-step until likelihood function converges. For each setting, we initialize multiple runs (see Methods) and keep the sequence with the highest converged likelihood.

##### 1.3 Approximating the confidence interval of latent curves: Variational inferences

The EM algorithm provides an efficient way to maximize the likelihood, but we only get a point estimate of $\theta_i$ without uncertainty quantification. Since we are interested in how genetic risk changes over age, we would ideally want to get the posterior of the latent risk curves over age. As discussed in previous sections, any sampling method will cause the "label switching"" problem, which will be hard to deal with, so we choose an approximation method. Here we apply "one step" of a Variational Bayes update to approximate the posterior distribution of the latent curves, the validity of which is discussed later in this section. We assume a factorized Variational distribution $q(z, \theta) = q(z)q(\theta)$. The full data distribution is as follows:

$$P(\hat{\beta}, \theta, \mathbf{Z}) = \prod_{i=1}^{K} \prod_{j=1}^{S} P(\hat{\beta}_j | \theta_i, \mathbf{Z}) P(\mathbf{Z}) P(\theta) = \prod_{i=1}^{K} \left\{ \mathcal{N}(\theta_i | 0, \Sigma_0) \prod_{j=1}^{S} \{\pi_i \mathcal{N}(\hat{\beta}_j | X\theta_i, \Sigma_j)\}^{z_{ji}} \right\} \tag{10}$$

In order to get the posterior distribution of the latent profiles, we derive the update of the variational distribution $q^*(\theta_i)$ as follows:

$$\begin{aligned} \ln q^*(\theta_i) =& E_z[\ln P(\hat{\beta}|\mathbf{Z}, \theta_i) + \ln P(\theta_i)] + const \\ =& -\frac{1}{2} \theta_i^T (X^T \{\sum_{j=1}^{S} E[z_{ji}] \Sigma_j^{-1}\} X + \Sigma_0^{-1}) \theta_i \\ &+ \theta_i^T X^T \sum_{j=1}^{S} E[z_{ji}] \Sigma_j^{-1} \hat{\beta}_j + const, \end{aligned} \tag{11}$$

$$
\begin{aligned}
q^*(\theta_i) &= \mathcal{N}(\theta_i|\mathcal{A}_i^{-1}\mathbf{b}_i, \mathcal{A}_i^{-1}) \\
\mathcal{A}_i &= X^T\{\sum_{j=1}^{S} E[z_{ji}]\Sigma_j^{-1}\}X + \Sigma_0^{-1} \\
\mathbf{b}_i &= X^T \sum_{j=1}^{S} E[z_{ji}]\Sigma_j^{-1}\hat{\beta}_j.
\end{aligned}
$$

Since $\theta_i$ is Gaussian, the latent profile $X\theta_i$ is also Gaussian (cubic spline is a linear transformation of $\theta_i$) with covariance matrix $X\mathcal{A}_i^{-1}X^T$.

We note that full Variational Bayes inference could potentially be performed by estimating $q^*(z_{ji})$ and $\pi_i$ by maximizing the Variational lower bound of the marginal likelihood.[3, p. 484] However, for Variational inference we use a factorized distribution $q(z,\theta) = q(z)q(\theta)$ to approximate the true posterior distribution of $P(\mathbf{Z},\theta|\hat{\beta})$, which can not be factorized. So, directly using Variational Bayes to estimate the mean of $\theta$ that maximizes Equation 5 is not as good as EM if we want an unbiased estimate. We therefore use EM to estimate the mean and the Variational Bayes approach to estimate the credible interval.

It is interesting to notice in Equation 11, the only expectation we took was over $z_{ji}$, which is also estimated in the EM algorithm's E-step, as in Equation 9. We now show how directly plugging in the expectation from EM will give a good approximation for the posterior distribution of latent curves.

Recall that the EM algorithm maximizes the marginal likelihood by iteratively maximizing the lower bound $L_{EM}(q,\theta)$ with respect to $q$ and $\theta$. The estimated value after converging can be expressed as:

$$
\begin{aligned}
(\hat{q}_{EM}(\mathbf{Z}), \hat{\theta}_{EM}) &= \arg\max_{q,\theta} L_{EM}(q(\mathbf{Z}),\theta) \\
&= \arg\max_{q,\theta} \sum_{z\in\mathbf{Z}} q(z) \ln \frac{P(\hat{\beta}, z|\theta)P(\theta)}{q(z)}
\end{aligned} \tag{12}
$$

The variational Bayes method also maximizes a lower bound for the marginal likelihood $L_{VB}(q)$, which can be written as:

$$
\begin{aligned}
(\hat{q}_{VB}(\theta), \hat{q}_{VB}(\mathbf{Z})) &= \arg\max_{q} L_{VB}(q) \\
&= \arg\max_{q} \int_\theta \sum_{\mathbf{Z}} q(z)q(\theta) \ln \frac{P(\hat{\beta}, z|\theta)P(\theta)}{q(z)q(\theta)} \\
&= \arg\max_{q} \int_\theta q(\theta) \sum_{\mathbf{Z}} q(z) \ln \frac{P(\hat{\beta}, z|\theta)P(\theta)}{q(z)} + \int_\theta q(\theta) \ln \frac{1}{q(\theta)} \\
&= \arg\max_{q} \int_\theta q(\theta) L_{EM}(q(\mathbf{Z}),\theta) - \int_\theta q(\theta) \ln q(\theta).
\end{aligned} \tag{13}
$$

In Equation 12, if we plug in the $\hat{q}_{EM}(\mathbf{Z})$ estimated via EM, we see that the lower bound $L_{EM}(\hat{q}_{EM}(\mathbf{Z}), \theta) = g(\theta)$ is a function of $\theta$, which is maximized at $\hat{\theta}_{EM}$. We note that the Variational lower bound in Equation 13 is the negative $KL(q||g)$, which is maximized when $q = g$. Comparing Equation 11 and Equation 12, we can see that $g(\theta)$ is estimated as $q^*(\theta)$ on the right hand side of Equation 11, with respect to $\hat{q}_{EM}(\mathbf{Z})$. Therefore, one step of Variational approximation of $q^*(\theta)$ is sufficient to provide an approximate posterior distribution for the latent curve for each cluster, which has the mode that maximizes the marginal likelihood (as in the EM algorithm).

### 2 The effect of unobserved factors on risk effect estimation

#### 2.1 Independent unobserved effects: frailty

Disease is caused by many risk factors, including those that are not observed. The unobserved confounders can cause biases for effect size estimation [4]. Here, we describe how unmeasured confounders affect risk effect size estimation, even when the confounders are independent of the focal variant of interest. We also provide methods to account for these effects through estimating frailty parameters from incidence rates of disease over age.

##### 2.1.1 Parametric modelling of effect size in the presence of frailty

Assuming we have a proportional hazards risk for disease incidence, with an unobserved risk background $u$, the hazard rate is given by

$$h(t|x, u) = h_0(t)e^{\beta x}u,$$

where $h(t|x, u)$ is the conditional hazard rate, $x$ is the focus genetic variant of interest. $u$ is the unobserved background risk effect (called frailty in epidemiology). Since $u$ is unobserved, instead of estimating the effect size $\beta$ and baseline risk $h_0(t)$ under the true model specification, we can only estimate the marginal effect size $\beta^*$ and baseline hazard $h_0^*(t)$ under a mis-specified model:

$$h(t|x) = h_0^*(t)e^{\beta^* x}. \tag{14}$$

Suppose we have a parametric form for $u$, we can then work out the marginal distribution for $\beta^*$ and $h_0^*(t)$ directly. For simplicity, here we assume that $u$ has a Gamma distribution [5]: $u \sim F(u) = Gamma(u|shape = a, scale = \theta)$. The marginal hazard ratio $h(t|x)$ can be computed as follows:

$$
\begin{aligned}
h(t|x) =& \frac{f(t|x)}{\mathcal{F}(t|x)} \\
=& \frac{\int_{\mathcal{U}} f(t|x,u)dF(u)}{\int_{\mathcal{U}} \mathcal{F}(t|x,u)dF(u)} \\
=& \frac{\int_{\mathcal{U}} h_0(t)e^{\beta x} u \exp(-ue^{\beta x}\int_0^t h_0(t)dt)dF(u)}{\int_{\mathcal{U}} \exp(-ue^{\beta x}\int_0^t h_0(t)dt)dF(u)} \\
=& h_0(t)e^{\beta x}\frac{\int_{\mathcal{U}} u^a \exp(-(\Lambda(t)e^{\beta x}+\frac{1}{\theta})u)\,du}{\int_{\mathcal{U}} u^{a-1}\exp(-(\Lambda(t)e^{\beta x}+\frac{1}{\theta})u)\,du} \\
=& h_0(t)e^{\beta x}\frac{\Gamma(a+1)}{\Gamma(a)}\frac{1}{\Lambda(t)e^{\beta x}+\frac{1}{\theta}} \\
=& h_0(t)e^{\beta x+\ln a-\ln(\frac{1}{\theta}+\Lambda(t)e^{\beta x})},
\end{aligned}
\tag{15}
$$

where $\Lambda(t) = \int_0^t h_0(t)dt$. Suppose $x \in \{0,1\}$ is a binary variable, then by comparing Equation 14 and Equation 15 we have:

$$\beta^* = \beta + \ln\frac{1+\theta\Lambda(t)}{1+\theta\Lambda(t)\exp(\beta)}.$$

Since most of the diseases have a low incidence rate in the population and the genetic risk sizes are relatively small, we can assume both $\Lambda(t)$ and $\beta$ to be small too. This leads to an approximation for $\beta^*$:

$$\beta^* \approx \beta(1-\theta\Lambda(t)), \tag{16}$$

which indicates that as age increases, we tend to increasingly underestimate the true risk. To account for the frailty effect in the clustering of curves, we change the intercept base in Equation 1 to be $(1-\theta\Lambda(t))$, and set the degrees of freedom of the model to be 1, which will infer a latent curve with an age dependency computed from the frailty model. The likelihood is then computed by plugging in the latent curve in Equation 4.

##### 2.1.2 Inferring frailty parameters from the population incidence rate

We describe an approach to fitting a parametric distribution $F(u)$) for frailty, with the constraint that the expectation, $E[u]$, equals one [5]. We use $u$ to represent the disease hazard heterogeneity, and assume that the baseline risk $h_0(t)$ increases monotonically for each individual. The net impact of frailty is to cause individuals with larger frailty $u$ to acquire the disease earlier, causing the incidence over the population to bend or even decrease for older age groups. We can fit a parametric hazard model with frailty to empirical incidence rates over age by assuming following parametric form of $h_0(t)$ and $F(u)$:

$$h(t|u) = u \cdot \gamma t^k$$

$$u \sim \mathcal{G}(shape = a, scale = \theta)\ s.t.\ E[u] = 1,\ var[u] = \theta.$$

Here we adopted a polynomial baseline hazard which represents the multi-stage nature of disease and a Gamma distribution for $F(u)$ [5]. We then compute the survival function:

$$\begin{aligned}
\mathcal{F}(t) &= \int_{\mathcal{U}} \exp(-u \int_0^t \gamma s^k ds) dF(u) \\
&= \int_{\mathcal{U}} \exp(-u \frac{\gamma t^{k+1}}{k+1}) \frac{\frac{1}{\theta^a}}{\Gamma(a)} e^{-\frac{u}{\theta}} u^{a-1} du \\
&= \frac{1}{(1 + \theta\gamma \frac{t^{k+1}}{k+1})^a}.
\end{aligned} \tag{17}$$

Considering $E[u] = a\theta = 1$, the hazard over the population is then computed as:

$$h(t) = -\frac{\mathcal{F}'(t)}{\mathcal{F}(t)} = \frac{\gamma t^k}{1 + \theta\gamma \frac{t^{k+1}}{k+1}}.$$

We then computed the empirical hazard rate $\hat{h}(t)$ from age 45 to age 70 in the UK Biobank cohort and subtracted the intercept to match the parametric form. The disease incidence after 70 years old is low and its analysis is complicated by competing health related events, giving an empirical hazard rate rate in the population that drops rapidly to zero. Therefore, we decided to only fit the model until age 70. The parameter $k, b, \gamma$ is then computed through the following least squares optimization:

$$argmin_{k,b,\gamma} \sum_t (\hat{h}(t) - h(t))^2$$

.

Examples of the fitted hazard rates using the frailty model are shown in Supplementary Figure 6, we then plug the fitted values $k, b, \gamma$ into Equation 16 to obtain the frailty correction term. We can then fit a genetic profile with frailty, as described in the Methods.

#### 2.2 Dependent unobserved effects: GxE and GxG interactions

Here, we show how interactions can induce the decreasing profile of genetic risk that we observe. Interactions between genetic risk factor and other risk factors, either observed or unobserved, can be modeled as an individual-level risk effect that is centered on the population level marginal effect size. Intuitively, if a risk allele affects individuals differently (because of the interactions), those individuals with the higher risk will tend to get the disease earlier, while those with lower risk will tend to have a later onset. When estimating risk effects at a particular age, we are essentially estimating the effect size from individuals whose disease onset is within that age group. Thus, we observe the decreasing risk over age.

To illustrate how such interactions can lead to a decreasing effect size (over age) for genetic risk factors, we consider a proportional hazard model with a single covariate. Following the model specification in Section 2.1, we use $\beta^*$ to represent the marginal effect size under a mis-specified model:

$$h(t) = h_0(t)\exp(\beta^* x), \tag{18}$$

where $h_0(t)$ is a positive baseline hazard and $x$ is the covariate. Assuming the effect size $\beta$ interacts with the environment or other genetic factors, we further assume that the effect size for each individual is generated from a positive defined probability distribution:

$$\beta \sim \mathcal{G}(\beta).$$

The survival function $\mathcal{F}(t)$ can be computed by integrating out $\beta$.

$$\begin{aligned}
\mathcal{F}(t) &= \int_{\beta\in\mathcal{G}} \mathcal{F}(t,\beta) \\
&= \int_{\beta\in\mathcal{G}} \exp(-e^{\beta x} * \int_0^t h_0(x)dx) \\
&= \int_{\beta\in\mathcal{G}} \exp(-e^{\beta x}\Lambda(t)).
\end{aligned} \tag{19}$$

As in Section 2.1, we use $\Lambda(t) = \int_0^t h_0(x)dx$. Assuming continuity in $\Lambda(t)$ and $\mathcal{G}(\beta)$, such that Fubini's theorem can be applied, we obtain the analytical form for the hazard rate:

$$\begin{aligned}
h(t) &= -\frac{\mathcal{F}'(t)}{\mathcal{F}(t)} \\
&= -\frac{\int_{\beta\in\mathcal{G}} h_0(t)e^{\beta x}\exp(-e^{\beta x}\Lambda(t))}{\int_{\beta\in\mathcal{G}} \exp(-e^{\beta x}\Lambda(t))} \\
&= h_0(t)\cdot\frac{\int_{\beta\in\mathcal{G}} e^{\beta x}\exp(-e^{\beta x}\Lambda(t))}{\int_{\beta\in\mathcal{G}} \exp(-e^{\beta x}\Lambda(t))}.
\end{aligned} \tag{20}$$

We first discuss the initial condition when $t \to 0$:

$$\lim_{t\to 0} h(t) = \lim_{t\to 0} h_0(t)\int_{\beta\in\mathcal{G}} e^{\beta x} = \lim_{t\to 0} h_0(t)E_{\mathcal{G}}[e^{\beta x}]$$

Under these conditions, the estimation of the marginal effect size $\lim_{t\to 0}\beta^*$ equals the expectation of $\beta$ under the distribution $\mathcal{G}$.

$$\lim_{t\to 0}\beta^* = \frac{\ln(E_{\mathcal{G}}[e^{\beta x}])}{x}.$$

Comparing Equation 18 and 20, we find:

$$\exp(\beta^* x) = \frac{\int_{\beta\in\mathcal{G}} e^{\beta x}\exp(-e^{\beta x}\Lambda(t))}{\int_{\beta\in\mathcal{G}}\exp(-e^{\beta x}\Lambda(t))}. \tag{21}$$

Since we have specified the baseline hazard rate $h_0(t) > 0$, we have $\Lambda(t)$ increasing monotonically with $t$. Therefore, we can take the derivative of Equation 21 with respect to $\Lambda(t)$ to analyse the gradient of $\beta^*$ at time $t$:

$$\begin{aligned}\frac{\partial \exp(\beta^* x)}{\partial \Lambda(t)} =& -\frac{1}{(\int_{\beta\in\mathcal{G}}\exp(-e^{\beta x}\Lambda(t)))^2}\cdot \\ &\big[-\big(\int_{\beta\in\mathcal{G}}(e^{\beta x})^2\exp(-e^{\beta x}\Lambda(t)) * \int_{\beta\in\mathcal{G}}\exp(-e^{\beta x}\Lambda(t))\big)+ \\ &\big(\int_{\beta\in\mathcal{G}} e^{\beta x}\exp(-e^{\beta x}\Lambda(t))\big)^2\big] < 0,\end{aligned} \tag{22}$$

Where we make use of the Cauchy-Schwarz inequality, as

$$e^{\beta x}\exp(-\frac{1}{2}e^{\beta x}\Lambda(t)) > 0,\ \exp(-\frac{1}{2}e^{\beta x}\Lambda(t)) > 0.$$

. Comparing Equations 18, 20, 21, and 22, we conclude that the estimated marginal effect size $\beta^*$ will be underestimated as $t$ increases. This result could be extended to a general case of almost any probability distribution for $\mathcal{G}(\beta)$. Many different scenario could be fitted into this framework, all of which will lead to a decreasing genetic risk profile with age.
